# A versatile Tn7 transposon-based bioluminescence tagging tool for quantitative and spatial detection of bacteria in plants

**DOI:** 10.1101/2021.02.11.430857

**Authors:** Ayumi Matsumoto, Titus Schlüter, Katharina Melkonian, Atsushi Takeda, Hirofumi Nakagami, Akira Mine

## Abstract

Investigation of plant-bacteria interactions requires quantification of *in planta* bacterial titers by means of colony counting assays. However, colony counting assays are cumbersome and time-consuming, and are unable to detect spatial patterns of bacterial colonization in plants. Here, to overcome these shortcomings, we devised a broadly applicable genetic engineering tool for bioluminescence-based quantitative and spatial detection of bacteria in plants. We developed plasmid vectors that have broad host ranges and enable Tn*7* transposon-mediated integration of the *luxCDABE* luciferase operon into a specific genomic location ubiquitously found across bacterial phyla. These vectors allowed for generation of bioluminescent transformants of various plant pathogenic bacteria belonging to the genera *Pseudomonas, Rhizobium* (*Agrobacterium*), and *Ralstonia*. The bioluminescent transformant of *Pseudomonas syringae* pv. *tomato* DC3000 (*Pto*-lux) was as virulent in *Arabidopsis thaliana* as its parental strain. Direct luminescence measurements of *Pto*-lux-inoculated plant tissues reported bacterial titers in *A. thaliana, Solanum lycopersicum, Nicotiana benthamiana*, and *Marchantia polymorpha* as accurately as conventional colony counting assays. We further showed the utility of our vectors for converting the previously generated *Pto* derivatives to isogenic bioluminescent strains. Importantly, quantitative bioluminescence assays using these *Pto*-lux strains accurately reported the effects of plant immunity and bacterial effectors on bacterial growth with a dynamic range of 4 orders of magnitude. Moreover, macroscopic bioluminescence imaging illuminated spatial colonization patterns of the *Pto*-lux in/on inoculated plant tissues. Taken together, our vectors offer untapped opportunities for developing bioluminescence-based quantitative and spatial analysis of bacterial growth in a variety of plant-bacteria interactions.

**SIGNIFICANCE STATEMENT:** We developed broad-host-range plasmid vectors that integrate the luciferase operon, *luxCDABE*, into a specific genomic location ubiquitously found across bacterial phyla. Using these vectors, we established a high-throughput method for bioluminescence-based quantitative assays of *in planta* bacterial growth with a dynamic range of 4 orders of magnitude and visualized spatiotemporal patterns of bacterial colonization in/on inoculated plant tissues.

## INTRODUCTION

Plants are colonized by a multitude of microbes such as bacteria, fungi, and oomycete. Plant-colonizing microbes can be classified into pathogens, mutualists, and commensals that cause negative, positive, and neutral consequences to plant health, respectively. Bacterial pathogens provoke the plant immunity mediated by cell surface receptors but suppress the resulting defense responses by deploying a repertoire of effectors (Dou and Zhou, 2012), allowing them to massively proliferate inside plants and cause diseases. Plants can restrict pathogen growth when inducing another layer of plant immunity known as effector-triggered immunity (ETI) through recognition of specific effectors by intracellular nucleotide-binding leucine-rich repeat receptors (NLRs) (Cui *et al*., 2015). Traditionally, *in planta* growth of bacteria is quantified by plating serial dilutions of tissue extracts on selective plates and counting the number of bacterial colonies appeared. This so-called colony counting assay has been instrumental to studies on interactions between plants and bacterial pathogens, providing molecular insights into the mechanisms of plant immunity as well as bacterial virulence. However, colony counting assays are cumbersome and time-consuming and demand skilled hands for reproducible results, which can be a bottleneck especially when high-throughput assays or large-scale quantitative data are required. Moreover, colony counting assays lose spatial information on bacterial colonization *in planta* because of tissue disruption in the sample preparation step.

Quantification of light emission from genetically engineered bioluminescent bacteria has been used as an alternative approach to estimate growth of pathogenic bacteria in infected animal and plant hosts (Fan *et al*., 2008, Howe *et al*., 2010, Cruz *et al*., 2014). For this purpose, bacterial luciferase operons such as *luxCDABE* of *Photorhabdus luminescens* and *Vibrio fischeri* are particularly useful. The *luxCDABE* operon expresses a heterodimeric luciferase encoded by *luxA* and *luxB*, which catalyzes the oxidation of reduced flavin mononucleotide (FMNH_2_) and a long-chain fatty aldehyde, resulting in the emission of light at a wavelength of 490 nm (Meighen, 1993). FMNH_2_ is continuously supplied in aerobically growing bacterial cells, while the long-chain fatty aldehyde is continuously regenerated by a fatty acid reductase complex encoded by *luxC, luxD*, and *luxE* (Meighen, 1993). Thus, metabolically-active bacteria expressing *luxCDABE* spontaneously emit light without supplying luciferase substrates.

The introduction of a plasmid vector carrying *luxCDABE* is an easy way to generate bioluminescent bacteria (Hikichi *et al*., 1998, Hikichi *et al*., 1999). However, bioluminescence emitted by such bacteria is influenced by plasmid copy numbers and is unstable without antibiotic selection. To circumvent these problems, stable chromosomal integration of *luxCDABE* has been employed (Tsuge *et al*., 1999, Fan *et al*., 2008, Howe *et al*., 2010, Cruz *et al*., 2014). For instance, Fan *et al*. (2008) used an artificial transposon that induces random transposition of *luxCDABE* to generate bioluminescent *Pseudomonas syringae* pv. *tomato* DC3000 (*Pto*) and *P. syringae* pv. *maculicola*. They demonstrated that the bioluminescence emitted by these bacteria can be used as a proxy to quantify their growth in *Arabidopsis thaliana* leaves. However, the use of these bioluminescent *P. syringae* strains has been limited due to the lack of evidence that the random insertion of *luxCDABE* did not cause undesired gene disruption. Determining genomic location of randomly inserted *luxCDABE* takes time but is essential if further genetic analysis of bacterial genes is planned.

Among transposable elements, the transposon Tn*7* is characterized by its integration mode. Transposition of Tn*7* occurs at a specific and neutral site known as Tn*7* attachment (attTn*7*) in a specific orientation at high frequency. AttTn*7* is located downstream of the *glmS* gene encoding an essential glucosamine-fructose-6-phosphate aminotransferase (Peters and Craig, 2001) and is found across bacterial phyla including *Proteobacteria, Firmicutes*, and *Bacteroidetes* (Wiles *et al*., 2018). Although chromosomal tagging with *luxCDABE* at attTn*7* has been successfully used to monitor growth of animal pathogenic bacteria (Howe *et al*., 2010), its use in plant pathogenic bacteria has not yet been reported. A major reason for this is the lack of an efficient and broadly applicable tool for Tn*7*-mediated chromosomal tagging of plant-colonizing bacteria with *luxCDABE*.

Here, we addressed the above-mentioned issue by developing Tn*7*-based vectors for chromosomal integration of *Photorhabdus luminescens luxCDABE*. The broad applicability of these vectors was demonstrated by the successful bioluminescence tagging of various plant pathogenic bacteria belonging to the genera *Pseudomonas, Rhizobium* (*Agrobacterium*), and *Ralstonia*. We then provided the evidence that our bioluminescence assays with the bioluminescent *Pto* (*Pto*-lux) and its derivatives quantitatively evaluate the effects of plant immunity and bacterial virulence on bacterial growth in a wide range of plant species. Moreover, using *Pto*-lux, we visualized spatially and temporally variable patterns of bacterial colonization in/on inoculated tissues of several plant species.

## RESULTS

### Construction of pBJ vectors

*Pto* can infect crops such as tomato as well as model plants such as *A. thaliana* and the liverwort *Marchantia polymorpha*, and thus serves as a model bacterial pathogen to study molecular and evolutionary plant-bacterial pathogen interactions (Xin *et al*., 2018, Gimenez-Ibanez *et al*., 2019). Previously, a bioluminescent *Pto* was generated by transposon-mediated random insertion of *luxCDABE* into the genome and was used for quantification of bacterial growth in *A. thaliana* (Fan *et al*., 2008). However, the use of this bioluminescent *Pto* strain has been limited, as random insertion of *luxCDABE* may have caused undesired gene disruption. To circumvent this problem, we thought to employ pBEN276, which was successfully used for integrating *luxCDABE* into *Salmonella enterica* (Howe *et al*., 2010). However, our attempt to transform *Pto* with pBEN276 was not successful likely because the replicon of the plasmid is not compatible for *Pto*.

To enable Tn*7*-mediated chromosomal integration of *luxCDABE* into *Pto* and other plant-colonizing bacteria, we took advantage of the broad-host-range vectors pBBR1-MCS2, pBBR1-MCS5, and pLAFR3 (Staskawicz *et al*., 1987, Kovach *et al*., 1995). We constructed seven vectors designated as pBJ1 to pBJ7 with two different constitutive promoters for *luxCDABE* expression and with three different antibiotic resistance genes (Figure 1 and Table 1). Although we introduced pBJ vectors into bacteria by electroporation in the present study, triparental mating can be also used as all pBJ vectors possess *mob* gene. After introduction of pBJ vectors into bacteria, Tn*7*-mediated transposition of *luxCDABE* can be induced by arabinose.

**Table 1.** Features of pBJ vectors. pBJ vectors possess either of the constitutive promoters P_frr_ or P_kan_ for *luxCDABE* expression and have different broad-host-range replicons and antibiotic resistance genes to broaden their applicability. All pBJ vectors possess *mob* gene and therefore can be transferred via triparental mating.

| Vector name | Promoter | Replicon | Resistance | Vector backbone | Other features |
| --- | --- | --- | --- | --- | --- |
| pBJ1 | P<sub>frr</sub> | pBBR1 | Gentamycin | pBBR1-MCS5 | mob |
| pBJ2 | P<sub>kan</sub> | pBBR1 | Gentamycin | pBBR1-MCS5 | mob |
| pBJ3 | P<sub>frr</sub> | pBBR1 | Kanamycin | pBBR1-MCS2 | mob |
| pBJ4 | P<sub>kan</sub> | pBBR1 | Kanamycin | pBBR1-MCS2 | mob |
| pBJ5 | P<sub>frr</sub> | pBBR1, RK2 | Gentamycin | pBBR1-MCS5 | mob |
| pBJ6 | P<sub>kan</sub> | pBBR1, RK2 | Gentamycin | pBBR1-MCS5 | mob |
| pBJ7 | P<sub>kan</sub> | RK2 | Tetracyclin | pLAFR3 | mob |

**Figure 1.**
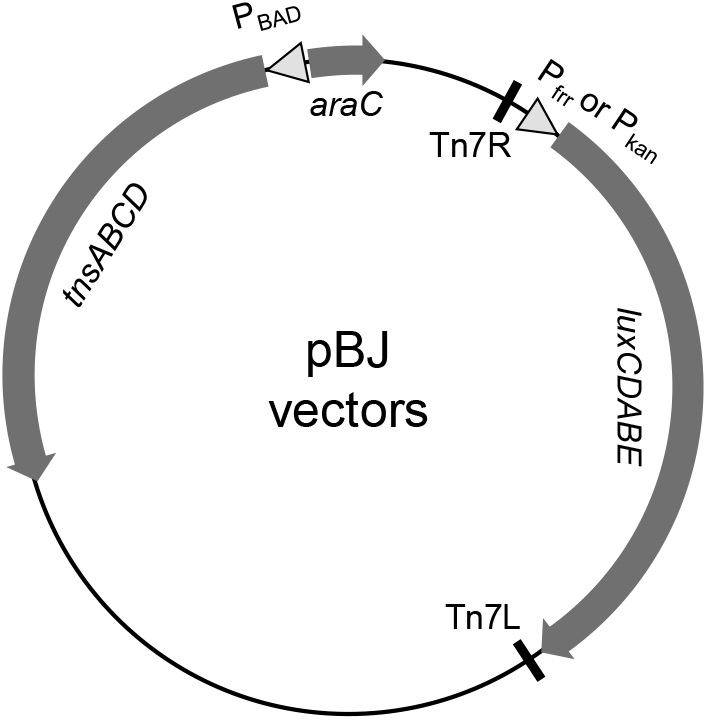
Schematic representation of pBJ vectors. *tnsABCD* are the genes required for transposition at attTn*7* and are controlled by an arabinose-inducible promoter (P_BAD_). The *luxCDABE* operon encoding luciferase is flanked by Tn*7* transposon arms Tn*7*L and Tn*7*R (indicated by vertical lines) and is controlled by either of the constitutive promoters P_frr_ or P_kan_. Detailed information for the features of pBJ vectors is available in Table 1.

### pBJ vectors enabled site-specific chromosomal integration of *luxCDABE* into a variety of plant pathogenic bacteria

Using pBJ vectors, we succeeded to generate bioluminescent transformants of *Pto* and *Agrobacterium tumefaciens* with available genome sequences, and Japanese isolates of *P. syringae* and *Ralstonia solacearum* without available genome sequences (Figure 2a and figure S1). Sequencing analysis confirmed that integration of *luxCDABE* occurs consistently at attTn*7* located 25 bp downstream of *glmS* in all transformed bacterial strains/species (Table S1). We did not observe growth defects of these bioluminescent bacteria *in vitro* (Figure 1A and figure S1), which is consistent with the generally accepted idea that attTn*7* is a neutral genomic site for bacterial fitness (Peters and Craig, 2001).

**Figure 2.**
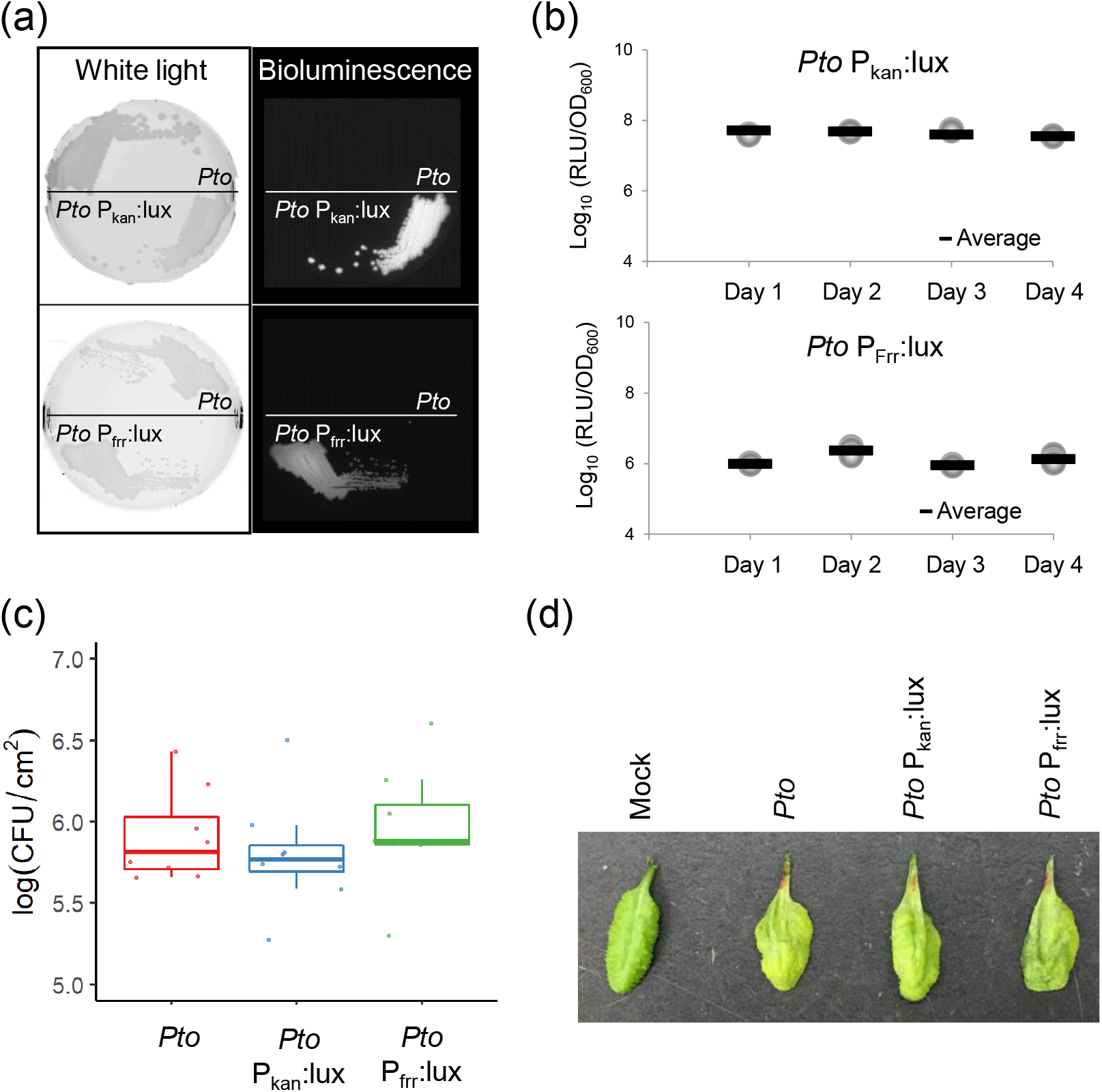
Characterization of *Pseudomonas syringae* pv. *tomato* DC3000 (*Pto*) with insertion of *luxCDABE* at attTn*7*. (a) *Pto* and *Pto* tagged with P_kan_- or P_frr_-driven *luxCDABE* (*Pto* P_kan_:lux or *Pto* P_frr_:lux) were observed under white light (left panels), and their bioluminescence was detected in the dark (right panels). *(b) Pto* P_kan_:lux or *Pto* P_frr_:lux were cultured in King’s B media without antibiotic selection for 4 days. Aliquots of overnight cultures were transferred to new media every day. Bioluminescence was quantified every day and normalized to bacterial density (OD_600_). (c)Growth of *Pto, Pto* P_kan_:lux and *Pto* P_frr_:lux (infiltrated at OD_600_ = 0.001) in leaves of *A. thaliana* Col-0 were determined at 3 days post inoculation (dpi) by standard plating assays. For each treatment, 8 biological replicates collected from different leaves were analyzed. (d) Disease symptom at 3 dpi of *A. thaliana* Col-0 leaves infiltrated with mock, *Pto, Pto* P_kan_:lux or *Pto* P_frr_:lux at OD_600_ = 0.01.

We compared activities of two different constitutive promoters, P_frr_ and P_kan_, used for *luxCDABE* expression. *Pto* with P_kan_-driven *luxCDABE* (*Pto* P_kan_:lux) showed stronger luminescence than *Pto* with P_frr_-driven *luxCDABE* (*Pto* P_frr_:lux) *in vitro* (Figures 2A and B). Both *Pto* derivatives maintained light emission for 4 days in non-selective *in vitro* cultures (Figure 2b), indicating that *luxCDABE* is stably maintained in the *Pto* genome. We then inoculated *A. thaliana* accession Col-0 with *Pto* P_kan_:lux and *Pto* P_frr_:lux, and measured bacterial titers at 3 days post inoculation (dpi) using conventional colony counting assays. *Pto* P_kan_:lux and *Pto* P_frr_:lux grew to a similar level to their parental wild-type *Pto* (Figure 2c). Disease symptoms caused by *Pto, Pto* P_kan_:lux, and *Pto* P_frr_:lux were comparable (Figure 2d). Taken together, we concluded that the stable integration of *luxCDABE* at attTn*7* does not affect *Pto* growth and virulence in *A. thaliana*.

### Bioluminescence accurately reported bacterial titers in various plant-*Pto* interactions

We next tested whether bioluminescence emitted by *Pto* P_kan_:lux and *Pto* P_frr_:lux can be used as a proxy to estimate bacterial titers *in planta. Pto* P_kan_:lux and *Pto* P_frr_:lux were syringe-infiltrated into leaves of *A. thaliana* Col-0 at varying doses. Leaf discs were excised from the bacteria-infiltrated leaves and placed in a light reflecting, white 96 well plate, followed by direct measurement of bioluminescence intensities (relative light units; RLUs) (Figure S2a). After bioluminescence measurement, the same leaf discs were subjected to conventional colony counting assays to determine bacterial titers (colony forming units; CFUs) (Figure S2b). We found that RLUs and CFUs show a strong positive correlation with a regression slope close to 1 (Figure 3a and figure S3a). Similar results were obtained with a Japanese isolate of *P. syringae* SUPP1331 with P_frr_-driven *luxCDABE* (Figure S3b). We noticed that bioluminescence emitted by *Pto* P_kan_:lux was approximately 10 times greater than that emitted by *Pto* P_frr_:lux in infected leaves (Figure 3a and figure S3a). Since higher bioluminescence emission increases the lower detection limit, we decided to employ *Pto* P_kan_:lux (hereafter referred to as *Pto*-lux) for further bioluminescence assays in this study.

**Figure 3.**
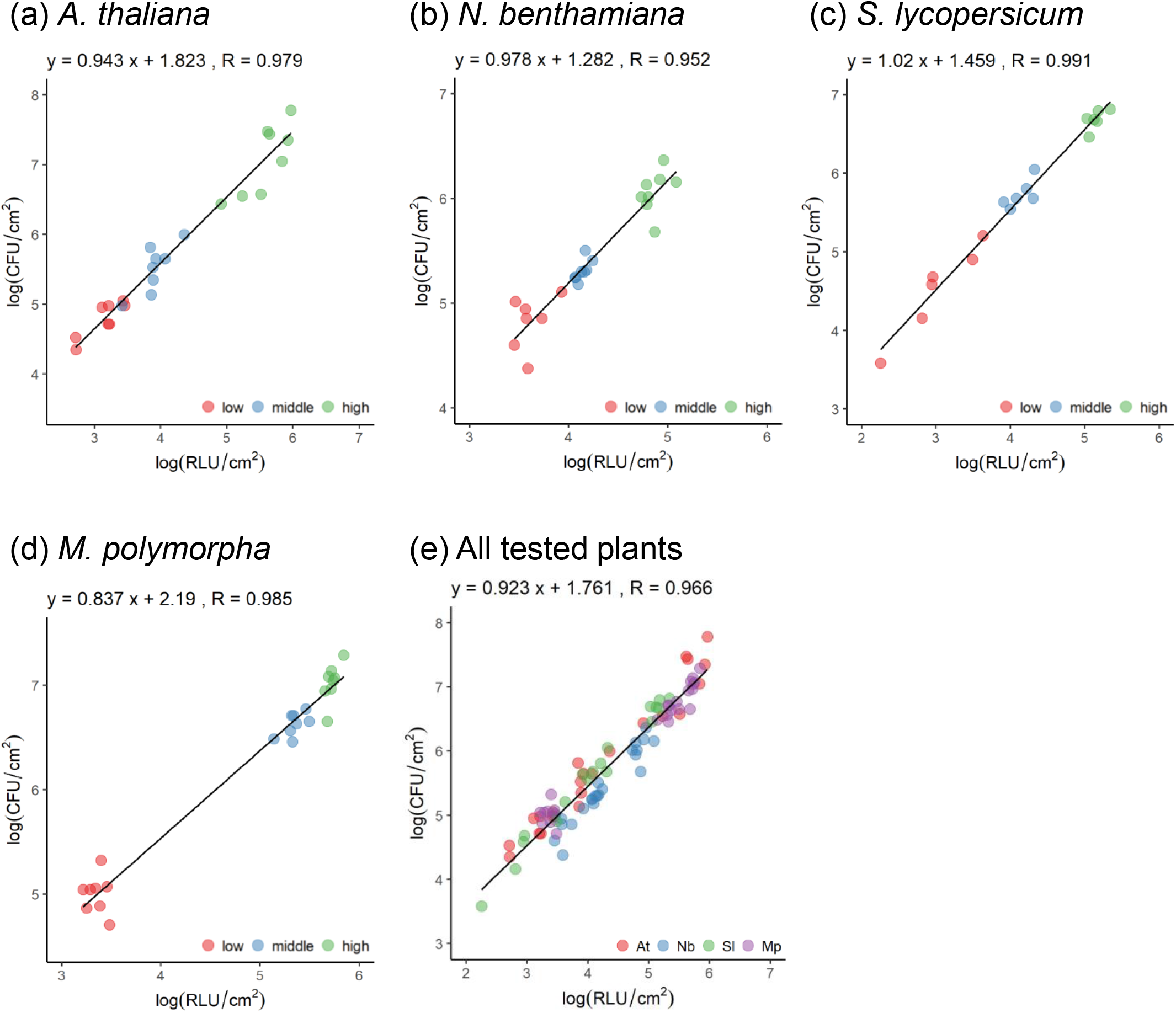
Bioluminescence accurately reports bacterial growth in various plant-*Pto* interactions. (a-d) *A. thaliana* (a), *Nicotiana benthamiana* (b), *Solanum lycopersicum* (c) and *Marchantia polymorpha* (d) were inoculated with *Pto* P_kan_:lux (*Pto*-lux hereafter) at varying doses (From low to high: OD_600_ = 0.0002, 0.001, 0.01 for *A. thaliana*; 0.00001, 0.00005, 0.0005 for *N. benthamiana*; 0.000002, 0.00001, 0.0001 for *S. lycopersicum*; 0.001, 0.01, 0.1 for *M. polymorpha*). Log_10_-transformed colony forming unit (CFU) and relative light unit (RLU) are plotted along with Y and X axes, respectively. Regression equations and Pearson’s correlation coefficients (R) are shown above the plots. (e) Similar correlation analysis was performed using the data from all the tested plants. At, *A. thaliana*. Nb, *N. benthamiana*. Sl, *S. lycopersicum*. Mp, *M. polymorpha*.

*N. benthamiana, S. lycopersicum*, and *M. polymorpha* have also been used in combination with *Pto* to explore molecular and evolutionary aspects of plant-bacterial pathogen interactions. *N. benthamiana* elicits ETI against *Pto* through recognition of HopQ1-1 effector (Wei *et al*., 2007). *S. lycopersicum* cv. M82 and *M. polymorpha* Takaragaike accessions, Tak-1 and Tak-2, are susceptible to *Pto* (Du *et al*., 2014, Gimenez-Ibanez *et al*., 2019). We tested whether RLU of *Pto*-lux can also be used as a proxy to measure *Pto* growth in these plant species. Leaves of *N. benthamiana* and *S. lycopersicum* cv. M82 were syringe-infiltrated with *Pto*-lux, while thalli of *M. polymorpha* Tak-1 was vacuum-infiltrated. The bacterial inoculation of *N. benthamiana* and *S. lycopersicum* and that of *M. polymorpha* were performed independently in two different laboratories. As shown in Figure 3, we found strong positive correlations between RLUs and CFUs and regression slopes close to 1 in all tested plant species. To our surprise, a strong positive correlation with a regression slope close to 1 was still evident when we plotted RLUs and CFUs obtained from different plant species all together (Figure 3e), even though the data was obtained in two laboratories using different experimental conditions. These results indicate that our quantitative bioluminescence assay is a highly reliable and robust measure of *in planta* bacterial titers.

### Development of the *Pto*-lux system for evaluation of the effects of plant immunity and bacterial effectors on bacterial growth in *A. thaliana*

We sought to make use of *Pto*-lux to recapitulate various aspects of the *A. thaliana*-*Pto* pathosystem. In *A. thaliana* Col-0, the recognition of AvrRpt2 and AvrRpm1 effectors by the corresponding NLRs, RPS2 and RPM1, respectively, triggers ETI to restrict *Pto* growth (Mackey *et al*., 2002, Axtell and Staskawicz, 2003, Mackey *et al*., 2003). *Pto*-lux was transformed with pLAFR3 carrying AvrRpt2 (*Pto*-lux AvrRpt2), AvrRpm1 (*Pto*-lux AvrRpm1), or empty pLAFR3 (*Pto*-lux EV). The resulting *Pto*-lux derivatives were used to inoculate *A. thaliana* Col-0 and *rpm1 rps2* plants. Both conventional colony counting assays and bioluminescence assays detected reduction in the growth of *Pto*-lux AvrRpt2 and *Pto*-lux AvrRpm1 compared to *Pto*-lux EV in Col-0 (Figure 4a and b). In *rpm1 rps2*, the growth of *Pto*-lux AvrRpt2 and *Pto*-lux AvrRpm1 was slightly higher than that of *Pto*-lux EV (Figure 4a and b). This likely reflects the virulence effects of AvrRpt2 and AvrRpm1 effectors in the absence of the corresponding receptors (Tsuda *et al*., 2009a). Importantly, we observed a strong positive correlation between RLUs and CFUs with a regression slope close to 1 (Figure 4c). These results demonstrated that our bioluminescence assays can quantitatively evaluate the effects of ETI and bacterial effectors on bacterial growth.

**Figure 4.**
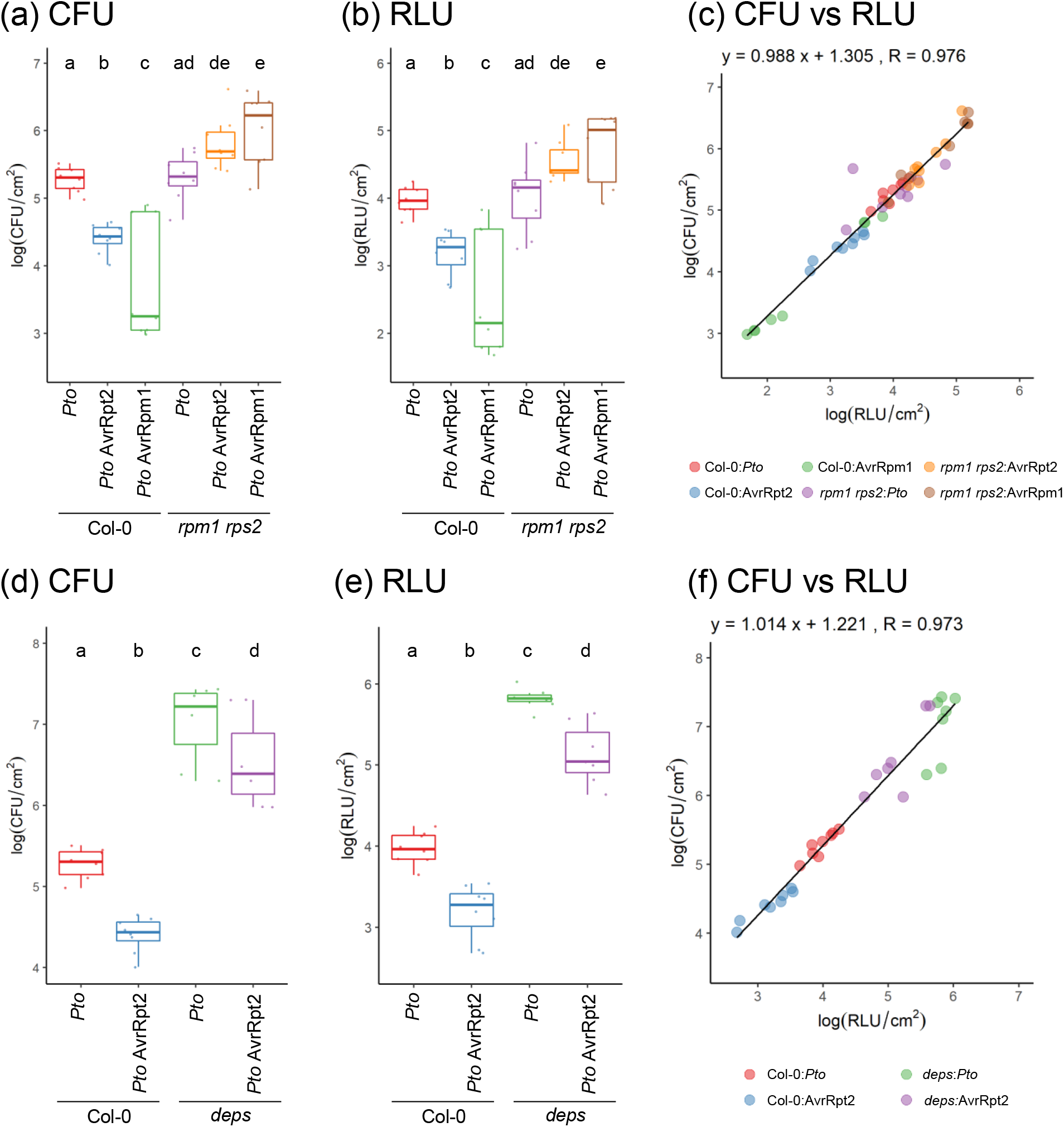
Bioluminescence assays accurately evaluate the effects of plant immunity on *Pto* growth in *A. thaliana*. (a-c) Leaves of Col-0 and *rpm1 rps2* plants were infiltrated with *Pto* P_kan_:lux (*Pto*-lux hereafter)carrying empty vector, AvrRpt2 or AvrRpm1 at OD_600_ = 0.001. Log_10_-transformed CFU (a) and RLU (b) at 2 dpi were calculated from 8 biological replicates collected from different leaves and were plotted along with Y and X axes, respectively (c). (d-f) Leaves of Col-0 and *dde2 ein2 pad4 sid2* (*deps*) were infiltrated with *Pto-*lux carrying empty vector or AvrRpt2 at OD_600_ = 0.001. Log_10_-transformed CFU (d) and RLU (e) at 2 dpi were calculated from 8 biological replicates collected from different leaves and were plotted along with Y and X axes, respectively (f). (a, b, d and e) Statistically significant differences are indicated by different letters (adjusted P < 0.05).

We then inoculated Col-0 and *dde2 ein2 pad4 sid2* (*deps*) plants with *Pto*-lux EV and *Pto*-lux AvrRpt2. The *deps* quadruple mutant is defective in defense signaling mediated by the phytohormones ethylene, jasmonic acid, and salicylic acid, and is therefore highly susceptible to *Pto* and *Pto* AvrRpt2 (Tsuda *et al*., 2009a). Both colony counting assays and bioluminescence assays detected approximately 100-fold increase in the growth of *Pto*-lux EV and *Pto*-lux AvrRpt2 in *deps* compared to those in Col-0 (Figure 4d and e). Measured CFUs and RLUs showed a strong positive correlation and a regression slope close to 1 (Figure 4f). The results obtained using the virulent and avirulent *Pto*-lux derivatives in combination with *A. thaliana* Col-0 and *deps* plants collectively indicated that our bioluminescence assays have a dynamic range of 10^3^ to 10^7^ CFU/cm^2^, which can mostly cover the range of *Pto* growth in resistant and susceptible genotypes of *A. thaliana*.

Previous studies have reported a number of hypovirulent *Pto* mutants such as *Pto* hrcC^-^ and *Pto* Δ*AvrPto*Δ*AvrPtoB. Pto* hrcC^-^ is deficient in type III secretion system required for effector delivery (Yuan and He, 1996), whereas *Pto* Δ*AvrPto*Δ*AvrPtoB* lacks two functionally redundant effectors AvrPto and AvrPtoB (Yuan and He, 1996, Lin and Martin, 2005, He *et al*., 2006). We introduced *luxCDABE* to *Pto* hrcC^-^ and *Pto* Δ*AvrPto* Δ*AvrPtoB* (*Pto*-lux hrcC^-^ and *Pto*-lux Δ*AvrPto* Δ*AvrPtoB*) at attTn*7* by the use of pBJ2

(Figure S4). It is worth noting that the resulting bioluminescent strains can be considered isogenic to *Pto*-lux, as all strains carry *luxCDABE* at the same genomic location (Table S1). Colony counting assays and bioluminescence assays similarly showed that *Pto*-lux hrcC^-^ and *Pto*-lux Δ*AvrPto* Δ*AvrPtoB* grew less compared to *Pto*-lux in Col-0 (Figures 5a and b), which were further supported by a strong positive correlation between CFUs and RLUs with a regression slope close to 1 (Figure 5c). These results emphasize that pBJ vectors make it possible to generate isogenic bioluminescent strains of the previously generated *Pto* mutants for comparative bioluminescence-based growth assays.

**Figure 5.**
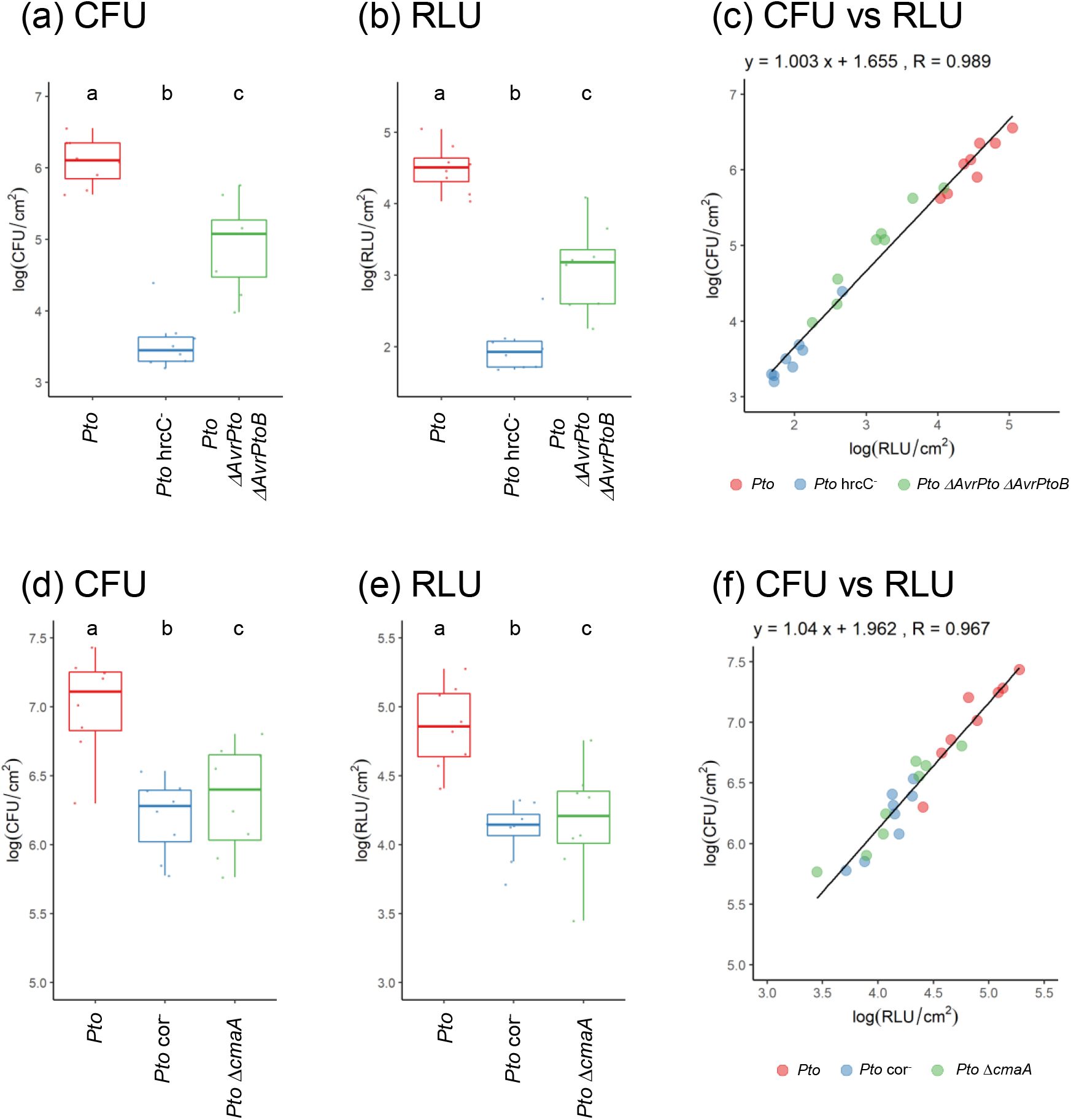
Bioluminescence assays accurately evaluate the effects of bacterial virulence factors on *Pto* growth in *A. thaliana*. (a-c) Leaves of Col-0 were infiltrated with *Pto-*lux, *Pto*-lux *hrcC-* or *Pto*-lux *ΔAvrPto ΔAvrPtoB* at OD_600_ = 0.0005. Log_10_-transformed CFU (a) and RLU (b) at 2 dpi were calculated from 8 biological replicates collected from different leaves of 3 plants and were plotted along with Y and X axes, respectively (c). (d-f) *A. thaliana* Col-0 plants were spray-inoculated with *Pto-*lux, *Pto*-lux cor- and *Pto*-lux *ΔcmaA* at OD_600_ = 0.1. Log_10_-transformed CFU (d) and RLU (e) at 3 dpi were calculated from 8 biological replicates, each of which was collected from 3 different leaves of a single plant, and were plotted along with Y and X axes, respectively (f). (a, b, d and e) Statistically significant differences are indicated by different letters (adjusted P < 0.05).

*Pto* produces the phytotoxin coronatine (COR) to re-open closed stomata, thereby gaining access into the apoplast for multiplication (Melotto *et al*., 2006). To investigate whether our bioluminescence assays can evaluate the function of COR, we used pBJ2 to generate a bioluminescent transformant of the COR-deficient strain *Pto* cor^-^ (Melotto *et al*., 2006) (*Pto*-lux cor^-^) (Figure S4). In addition, we used a suicide vector-based gene deletion to generate a *Pto*-lux mutant lacking *cmaA* gene required for COR production (Brooks *et al*., 2004) (*Pto*-lux *ΔcmaA*) (Figure S5). These bioluminescent strains were used for spray-inoculation of Col-0 plants. *Pto*-lux cor^-^ and *Pto*-lux *ΔcmaA* showed reduced growth compared to *Pto*-lux in both colony counting assays and bioluminescence assays where CFUs and RFUs showed a strong positive correlation and a regression slope close to 1 (Figure 5d-f). Thus, besides transformation of the existing *Pto* mutants by pBJ vectors, gene deletion approach with *Pto*-lux is also feasible for evaluating virulence functions of bacterial genes based on bioluminescence assays.

### Macroscopic bioluminescence imaging illuminated heterogenous colonization patterns of *Pto*-lux in infected leaves

Colony counting assays include a step of cell/tissue disruption and, therefore, lose spatial information on bacterial colonization patterns in infected tissues. We took advantages of *Pto*-lux to visualize the sites of bacterial colonization at a macroscopic level using a luminescence imager. Despite that suspensions of *Pto*-lux were infiltrated into whole areas of *A. thaliana* Col-0 leaves, we observed spotty bioluminescence signals in the infiltrated leaves (Figure 6a). Similar results were obtained in *S. lycopersicum* (Figure 6b). These results suggested that *Pto* multiplication is spatially variable in infected leaves.

**Figure 6.**
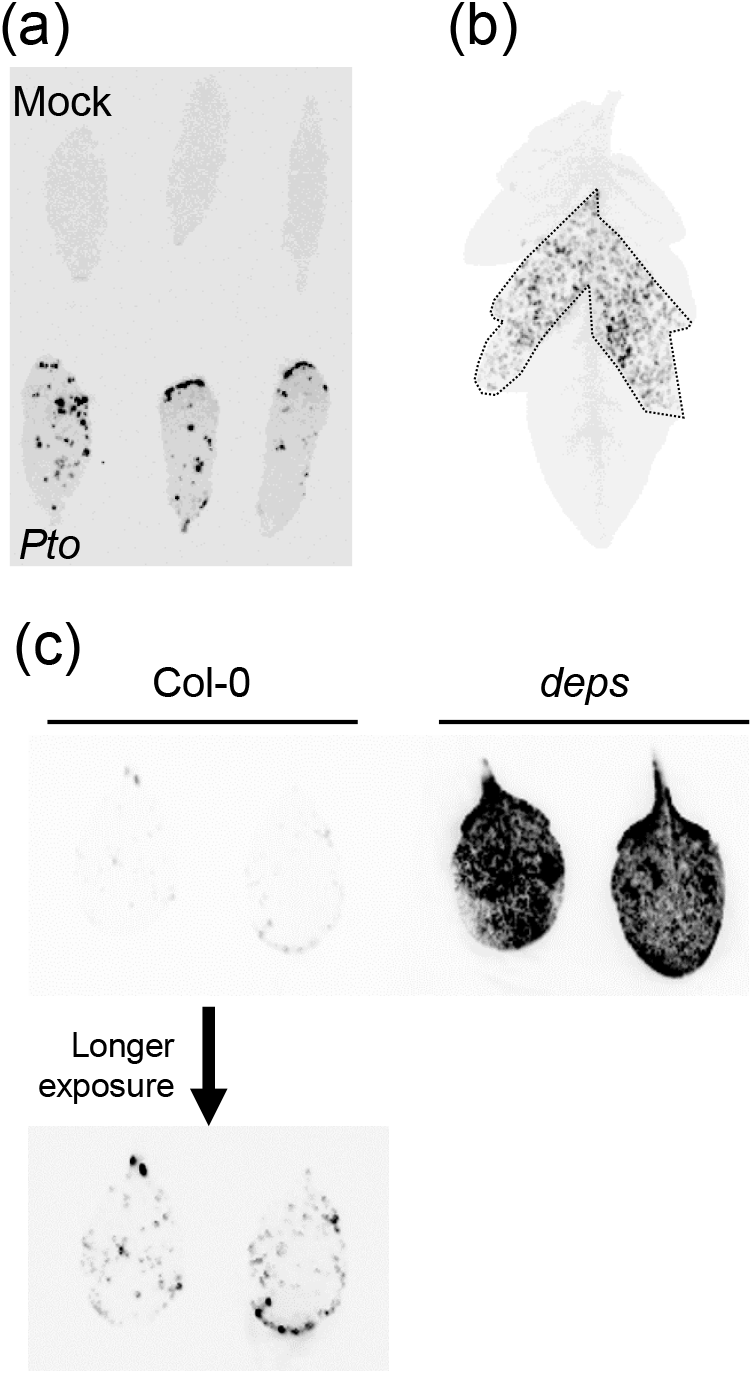
Bioluminescence illuminates heterogenous colonization of *Pto* in inoculated leaves. (a) Bioluminescence of *A. thaliana* Col-0 leaves was detected 2 days after infiltration with *Pto*-lux at OD_600_ = 0.001. (b) Bioluminescence of *S. lycopersicum* cv. M82 leaves was detected 2 days after infiltration with *Pto*-lux at OD_600_ = 0.0001. The dotted line surrounds the area where bacterial suspensions were infiltrated. (c) Bioluminescence of Col-0 and *deps* leaves was detected 2 days after infiltration with *Pto*-lux at OD_600_= 0.001.

Intriguingly, we observed bioluminescence signals of *Pto*-lux in nearly the entire leaves of immune-compromised *deps* plants of *A. thaliana* (Figure 6c), suggesting that plant immunity affects spatial colonization patterns of this bacterial pathogen. Collectively, these results emphasize the utility of our bioluminescence imaging approach for spatial analysis of plant-bacteria interactions.

### The growth of *Pto*-lux on *M. polymorpha* thalli

We further explored the utility of our *Pto*-lux system by investigating the interaction between *Pto* and the liverwort *M. polymorpha*, which is an emerging and attractive pathosystem for elucidating the evolution of molecular mechanisms underlying plant-bacteria interactions (Gimenez-Ibanez *et al*., 2019). Since growth and colonization patterns of *Pto* in *M. polymorpha* have not yet been well described, we monitored the growth of *Pto*-lux in *M. polymorpha* Tak-1 over time. When we inoculated with the bacterial suspension at OD_600_ = 0.01, *Pto*-lux growth at the basal regions of thalli saturated at 2 dpi and the bacterial titer was maintained at least up to 4 dpi in our experimental conditions (Figure 7a). Bioluminescence imaging revealed that *Pto*-lux preferentially colonize and grow at the basal region of thalli at early time points and later at the side edges of thalli (Figure 7b). The bioluminescence signals derived from the inoculum were observed outside the thalli at 0 dpi but diminished later than 1 dpi (Figure 7b). Disease symptoms associated with chlorosis were observed mainly at the basal region at later time points (Figure 7b). These temporal colonization patterns fit well to the bacterial titers that were measured at the basal region (Figure 7a). Thus, bioluminescence imaging with *Pto*-lux revealed temporal and spatial dynamic of *Pto*-*M. polymorpha* interactions.

**Figure 7.**
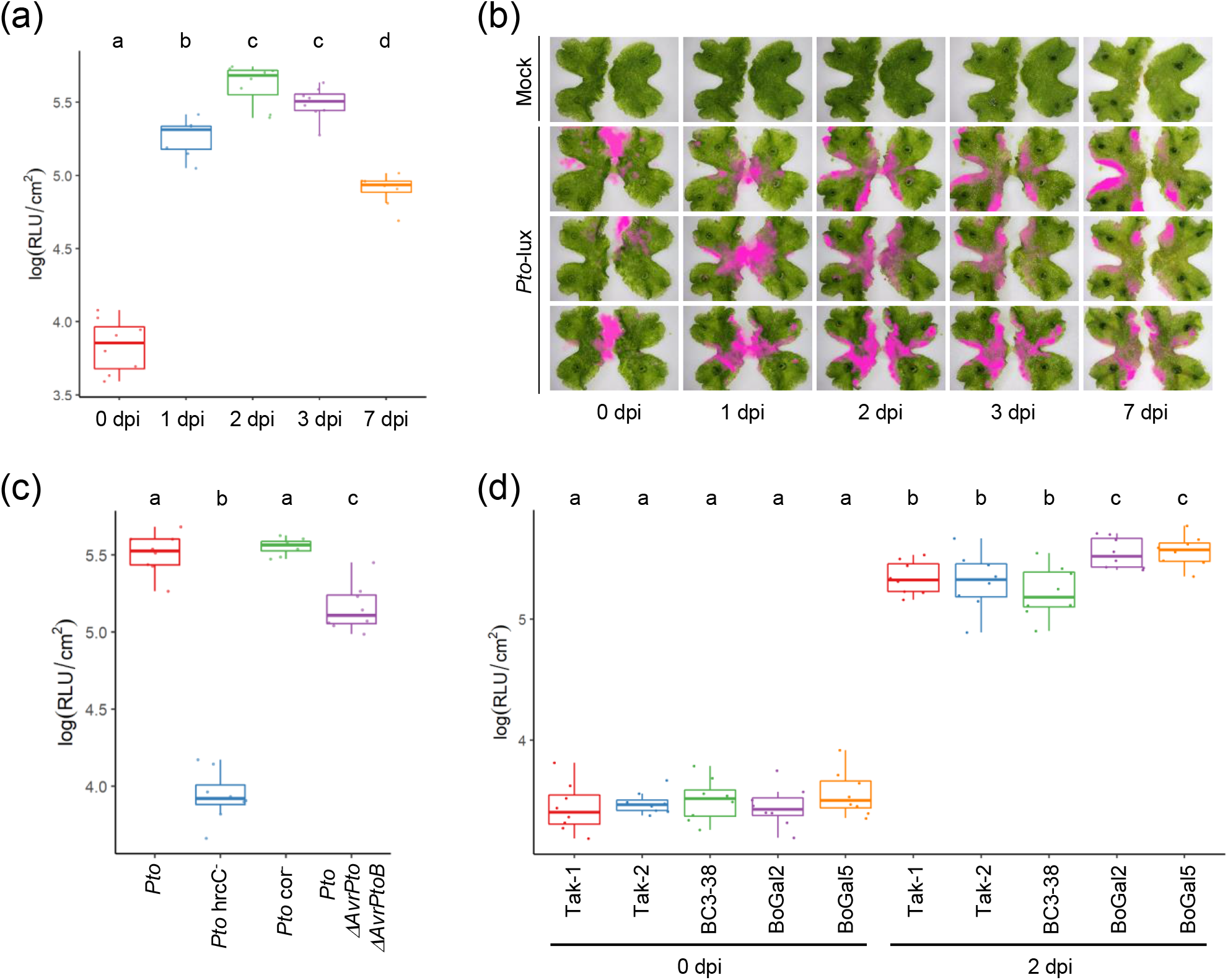
*Pto*-lux serves as a new tool for genetic dissection of *M. polymorpha-Pto* interactions. (a and b) Fourteen-days-old Tak-1 thalli were vacuum infiltrated with *Pto*-lux at OD_600_ = 0.01 and sampled at 0, 1, 2, 3, and 7 dpi. (a) Log_10_-transformed RLU were calculated from 8 biological replicates collected from thalli of different plants. (b) Bright field images were merged with pseudocolored bioluminescence images (pink). (c) Fourteen-days-old Tak-1 thalli were vacuum infiltrated with *Pto*-lux, *Pto*-lux hrcC-, *Pto*-lux *c*or- or *Pto*-lux Δ*AvrPto* Δ*AvrPtoB* at OD_600_ = 0.01. Log_10_-transformed RLU were calculated from 8 biological replicates collected from thalli of different plants. (d) Fourteen-days-old Tak-1, Tak-2, BC3-38, BoGa L2 and BoGa L5 thalli were vacuum infiltrated with *Pto*-lux at OD_600_ = 0.01. Log_10_-transformed RLU were calculated from 8 biological replicates collected from thalli of different plants. (a,c and d) Statistically significant differences are indicated by different letters (adjusted P < 0.05).

Virulence and infection of *Pto* to *M. polymorpha* depends on bacterial effectors, as shown by the use of *Pto* mutants, *Pto* hrcC^-^, *Pto* cor^-^, *Pto* Δ*AvrPto* Δ*AvrPtoB* (Gimenez-Ibanez *et al*., 2019). Therefore, we tested whether the reported observations can be recapitulated by using our *Pto*-lux system. Consistent with the previous study, our bioluminescence assay showed that a functional type III secretion system and the effectors AvrPto and AvrPtoB but not coronatine are required for full virulence of *Pto* against *M. polymorpha* (Figure 7c). This result assures that our bioluminescence assay is robust and compatible for evaluating functions of candidate effectors of bacteria in different plant models.

Currently, three *M. polymorpha* accessions, Tak-1 (male), Tak-2 (female), and BoGa (male and female lines), are broadly utilized as wild-type strains in molecular genetic studies (Okada *et al*., 2000, Ishizaki *et al*., 2008, Althoff *et al*., 2014, Buschmann *et al*., 2016). Spores produced by crossing Takaragaike accessions, Tak-1 and Tak-2, or Botanical Garden Osnabrück (BoGa) accessions have been used to generate mutant or transgenic plants and for forward genetic screening (Ishizaki *et al*., 2008, Ishizaki *et al*., 2013a, Ishizaki *et al*., 2013b, Sugano *et al*., 2014, Honkanen *et al*., 2016). However, these male and female lines very often display very distinct phenotypes. It is therefore very important to test whether these accessions differ in their levels of bacterial resistance. Tak-1 and Tak-2, and their backcrossed progeny, BC3-38, displayed the same level of resistance against *Pto*-lux (Figure 7d). BoGa-L5 (male) and BoGa-L2 (female) were comparably resistant. Thus, using *Pto*-lux, we provided the evidence that transgenic plants generated from spores produced by crossing Tak-1 and Tak-2 plants or BoGa-L5 and BoGa-L2 plants can be fairly compared to their parental plants in terms of bacterial resistance. Moreover, our bioluminescence assays revealed that BoGa-L5 and BoGa-L2 plants were slightly more susceptible compared to the Takaragaike accessions (Figure 7d). Finally, we made use of *Pto*-lux to compare different methods for bacterial inoculation in *M. polymorpha*. We first tested whether *Pto* can colonize *M. polymorpha* just by dipping thalli into the bacterial suspension without applying vacuum. As shown in Figure S6a, significant *Pto*-lux growth was observed after dip inoculation, although the bacterial titer at 2dpi was lower compared to vacuum infiltration. The usage of a filter paper on top of solid media for plant growth prior to inoculation sometimes affect thalli growth in an uncontrolled way, which is not observed for thalli grown directly on agar. Therefore, we tested whether thalli on agar plants without filter paper can be used for *Pto* infection assay (Figure S7a). We poured the bacterial suspension onto thalli grown on agar plates and vacuum infiltrated subsequently. The bacterial suspension was then poured off and thalli were cultured further on the same plates (Figure S7a). We found that the agar-based method is as feasible for measuring *Pto*-lux growth in *M. polymorpha* thalli as the above established filter-utilizing method (Figure S6b).

## DISCUSSION

Several studies reported the generation of bioluminescent bacteria chromosomally tagged with the *luxCDABE* luciferase operon and used them for studying plant-bacteria interactions (Tsuge *et al*., 1999; Fan *et al*., 2008; Cruz *et al*., 2014). However, these studies used genetic tools that are designed specifically to the bacterial strain used or that could cause undesired gene disruption by random insertion of the transgene. Thus, an efficient and broadly applicable tool for bioluminescence tagging is needed for researchers looking to manipulate their own bacterial isolates. In this study, we have developed pBJ vectors that can be used to readily generate bioluminescent bacteria by inducing Tn*7*-mediated chromosomal integration of *luxCDABE* into a specific and neutral site known as attTn*7*. It was shown that attTn*7* is conserved across bacterial phyla ranging from *Proteobacteria, Firmicutes*, to *Bacteroidetes* (Wiles *et al*., 2018). Selectable broad-host-range replicons and antibiotic resistance genes increase the applicability of pBJ vectors. Indeed, we generated bioluminescent transformants of various plant pathogenic bacteria of different genera including *Pseudomonas, Rhizobium* (*Agrobacterium*), and *Ralstonia*. Thus, pBJ vectors serve as versatile tools for bioluminescence tagging of a wide range of bacterial lineages.

Using pBJ vectors, we generated *Pto*-lux and showed its use for bioluminescence-based quantification of bacterial titers in a rapid and high-throughput manner. Unlike conventional colony counting assays, our bioluminescence assays do not require sample grinding, making and plating serial dilutions, and manual colony counting, thereby saving substantial time and human labors and reducing human errors (Figure S2). In addition, our bioluminescence assays can avoid release of plant metabolites during tissue disruption, which is inevitable in colony counting assays and may cause undesired effects on bacterial growth on plates. It may be important to pay attention to these effects, because increasing evidence suggests that various plant species accumulate defense metabolites in specialized cells such as trichomes and/or intracellular vesicles and organelle (e.g., oil body and vacuole) (Hatsugai *et al*., 2009, Huchelmann *et al*., 2017, Romani *et al*., 2020).

Our *Pto*-lux system has solved the limitations that are inherent to a similar system reported previously (Fan *et al*., 2008). Unlike random insertion of *luxCDABE* used by Fan *et al*., Tn*7*-mediated site-specific transposition in our system allowed for comparative analysis of the isogenic bioluminescent transformants of wildtype *Pto* and the previously generated *Pto* derivatives with altered virulence such as *Pto* AvrRpt2, *Pto* AvrRpm1, *Pto* hrcC^-^, *Pto* cor^-^, and *Pto* Δ*AvrPto* Δ*AvrPtoB* (Figures 4 and 5). These *Pto* derivatives have been extensively used to study molecular mechanisms of plant innate immunity and bacterial virulence (He *et al*., 2006, Kim *et al*., 2009, Cui *et al*., 2013, Geng *et al*., 2016, Mine *et al*., 2017, Mine *et al*., 2018). In addition, since the genomic location of *luxCDABE* in *Pto*-lux is clearly defined, it is straightforward to use homologous recombination techniques for targeted gene knockins and knockouts in *Pto*-lux. This is exemplified by the genetic analysis of the COR function with *Pto*-lux Δ*cmaA* (Figure 5d-e).

Macroscopic bioluminescence imaging revealed spatially variable growth of *Pto*-lux in various plant species. For instance, we showed that even when bacterial suspensions were infiltrated into the entire areas of *A. thaliana* leaves, the sites of *Pto*-lux colonization were locally confined (Figure 6a). In contrast, bioluminescence signals of *Pto*-lux were observed in the almost entire areas of the leaves of immune-compromised *deps* plants (Figure 6c). These results suggest that plant immunity affects spatial patterns of bacterial multiplication and/or colonization. We were also able to visualize spatial colonization patterns of *Pto*-lux in *M. polymorpha* Tak-1 thalli over time (Figure 7b), which generally support the reported observation that *Pto* growth was lower at meristematic apical notches compared to the basal regions of thalli (Gimenez-Ibanez *et al*., 2019). It is reasonable to speculate that reproductive organs such as apical notches and gemma cups are resistant to *Pto* infection to ensure survival of *M. polymorpha*. It would be an interesting future challenge to explore plant and bacterial factors that affect the spatially variable bacterial growth in plants.

Bioluminescence-based quantitative growth assays are very efficient and can be utilized to optimize experimental conditions for plant-bacteria interactions. As a remarkable example, we established an experimental condition in which Tak-1, Tak-2, and their progeny BC3-38 display the same level of bacterial resistance (Figure 7d), which assures fair genetic comparisons of transgenic or mutant plants to the parental Tak-1 and Tak-2. Our finding is in sharp contrast to Gimenez-Ibanez *et al*. which reported differential *Pto* growth on Tak-1 and Tak-2 (Gimenez-Ibanez *et al*., 2019). Major differences between the two studies are plant age and growth conditions. We used 14-day-old thalli grown under the continuous light, whereas Gimenez-Ibanez *et al*. used 2 to 4-week-old thalli grown under a long day condition (16h light/8h dark cycle). Since developmental differences between Tak-1 and Tak-2 become more prominent as they grow older, we speculate that usage of relatively young thalli and sampling from the defined basal region, where *Pto* preferentially colonize and grow, are key for fair comparison between the genotypes. Moreover, we found that the male and female lines of BoGa accessions are similarly resistant but slightly more susceptible than Takaragaike accessions to *Pto*-lux (Figure 7d). Thus, we argue that our experimental system with *Pto*-lux will facilitate genetic analysis of *M. polymorph*-*Pto* interactions.

Bacteria are causative agents of plant diseases and are, at the same time, major constituents of the collective assemblage of plant-colonizing microbes called the plant microbiota that affect plant health (Pfeilmeier *et al*., 2016, Hacquard *et al*., 2017). Thus, in addition to advancing our understanding of plant-bacterial pathogen interactions, it is becoming increasingly important to elucidate the mechanisms by which plants and bacterial members of the plant microbiota establish mutualistic relationships with each other. We anticipate that bioluminescence-based quantitative and spatial detection of bacteria enabled by pBJ vectors will serve as new tools to facilitate these rapidly growing research fields of plant-bacteria interactions.

## EXPERIMENTAL PROCEDURES

### Plant materials and growth conditions

*A. thaliana* accession Col-0 and its mutants *rpm1-3 rps2-101C* (Mackey *et al*., 2003) and *dde2-2 ein2-1 pad4-1 sid2-2* (Tsuda *et al*., 2009b), *N. benthamiana, S. lycopersicum* cv. M82, and *M. polymorpha* accessions Tak-1 (male), Tak-2 (female), BC3-38 (female), BoGa-L5 (male), and Boga-L2 (female) (Althoff *et al*., 2014, Bowman *et al*., 2017, Yamaoka *et al*., 2018) were used in this study. *A, thaliana* plants were grown in a climate chamber at 22°C with a 10 h light period and 60% relative humidity. *N. benthamiana* plants were grown in an air-conditioned room at 22°C with a 16 h light period. *S. lycopersicum* plants were grown in a climate chamber with 16 h light/8 h dark at 24°C during daylight/22°C at night and 60% relative humidity. *M. polymorpha* gemmae or thalli were grown in a walk-in growth chamber at 22°C under 50-60 μmol photons m^-2^s^-1^ continuous white LED light.

### Construction of pBJ vectors

The pBBR1-MCS5 derivative pJN105 was cut with *Xho*I and *Not*I. The larger fragment was treated with T4 polymerase followed by circularization with T4 ligase, yielding pJN105ΔpBAD. Two DNA fragments encoding *tnsABCD* or *luxCDABE* were amplified by PCR from pBEN276 (Howe *et al*, 2010) using the primer sets Pfrr_inside_F plus New_Tn7L_pJN_M13F or Pfrr_inside_R plus pBEN_Pamp_pJN_M13R, respectively. These DNA fragments were assembled using NEBuilder HiFi Assembly Cloning Kit (NEB, E2621) into *Apa*I-digested pJN105ΔpBAD, yielding pBJ1. Kanamycin promoter (P_kan_) was amplified from pBBR1-MCS2 using Pkan_pBJ_F2 and Pkan_pBJ_R2 and assembled into *Xho*I-digested pBJ1 to replace frr promoter (P_frr_), resulting in pBJ2. pBJ3 and pBJ4 were designed to confer kanamycin resistance instead of gentamicin resistance conferred by pBJ1/2. A DNA fragment was amplified from pBEN276 using Pfrr_inside_R plus pBEN_Pamp_pJN_M13R and another DNA fragment was amplified from pBJ1 using Pfrr_inside_F plus pBJ_dXhoI_Tn7R. These two fragments were then assembled into *Apa*I-digested pBBR1-MCS2, yielding pBJ3. pBJ4 was constructed by replacing P_frr_ in pBJ3 with P_kan_ as described above.

To broaden host range of pBJ vectors, a broad-host-range RK2 replicon was amplified from pFREE-RK2 using OriV_RK2_Fwd plus OriV_RK2_Rev and assembled into *Nae*I-digested pBJ1 and pBJ2 to produce pBJ5 and pBJ6, respectively. For the similar purpose, the broad-host-range cosmid vector pLAFR3 was cut with *Eco*RI and treated with T4 polymerase, followed by further digestion with *Pst*I. This linearized vector was used as a recipient for a *Nae*I-*Sbf*I restriction fragment of pBJ2 containing the *tnsABCD*-Pkan-*luxCDABE* cassette, yielding pBJ7.

Primers used for construction of pBJ vectors are listed in Table S2.

### Generation of bioluminescent bacteria

pBJ vectors were used to generate chromosomally *luxCDABE*-tagged *P. syringae, A. tumefaciens* and *R. solanacearum*. Antibiotics were used at the following concentrations: rifampin, 50 μg/ml; kanamycin, 50 μg/ml; gentamycin, 10 μg/ml, and tetracycline, 20 μg/ml. Single colonies of *P. syringae* pv. *tomato* DC3000 (*Pto*), *Pto* hrcC^-^, *Pto* cor^-^, *Pto* Δ*AvrPto* Δ*AvrPtoB*, and Japanese isolates of *P. syringae* SUPP1331 and SUPP1141 were inoculated into King’B (KB) media with appropriate antibiotics, grown overnight, washed twice with 300 mM sucrose at room temperature and suspended in 300 mM sucrose. The cells were stored at −80°C until use. The competent cells were transformed with pBJ1 (gentamicin resistance), pBJ2 (gentamicin resistance), pBJ4 (kanamycin resistance), pBJ5 (gentamicin resistance), pBJ6 (gentamicin resistance) or pBJ7 (tetracycline resistance) by electroporation using a Gene Pulser Xcell PC system (Bio-Rad). The electroporation condition was 2.5kV, 25µF and 200Ω. Bacteria were recovered for 2-4 hours at 28°C in KB media and then streaked onto KB plates with appropriate antibiotics and placed in an incubator at 28°C for 4-5 days. Bioluminescent colonies were detected using Fusion SOLO S (VILBER), inoculated into KB media containing 0.1% L-arabinose and cultured overnight at 28°C to induce transposition. The cultures were streaked onto KB plates with rifampicin and placed in an incubator at 28°C for 2-3 days until single colonies appeared. Bioluminescent colonies were patched onto KB plates with appropriate antibiotics to select gentamicin-, kanamycin- or tetracycline-sensitive colonies, confirming the absence of pBJ vectors. Insertion of *luxCDABE* into the attTn*7* sites of bacterial genomes were determined by sequencing. pLAFR3 cosmid vectors carrying AvrRpt2 or AvrRpm1 were introduced by electroporation into *Pto* chromosomally tagged with P_kan_-driven *luxCDABE* (*Pto*-lux) which had been generated with pBJ2.

A single colony of *A. tumefaciens* GV3101 was inoculated into LB medium containing rifampicin and gentamicin, grown at 28°C to logarithmic phase (OD_600_=0.8), collected by centrifugation and suspended in ice-chilled 20 mM CaCl_2_. The cells were stored at −80°C until use. The competent cells were mixed with pBJ3 or pBJ4 and flash-frozen in liquid nitrogen, followed by incubation at 37°C for 30 min. Bacteria were recovered in SOC media at 28°C for 2 hours and then streaked onto LB plates with rifampicin, gentamicin, and kanamycin. A single bioluminescent colony was then inoculated into LB medium containing 0.1% L-arabinose and cultured overnight at 28°C to induce transposition. The culture was streaked onto LB plates with rifampicin plus gentamycin and placed in an incubator at 28°C for 2 days until single colonies appeared. Bioluminescent colonies were patched onto LB plates with appropriate antibiotics to select kanamycin-sensitive colonies, confirming the absence of pBJ vectors. Chromosomal integration of *luxCDABE* was determined by sequencing.

A single colony of *R. solanacearum* OE1-1 (Kanda *et al*., 2003) was inoculated into PY medium, grown at 30°C to logarithmic phase (OD_600_=0.8), collected by centrifugation and suspended in ice-chilled 10% glycerol. The cells were stored at −80°C until use. The competent cells were transformed with pBJ7 by electroporation, the condition of which was 3kV, 25µF and 300Ω. Bacteria were recovered in SOC medium at 30°C for 2 hours and then streaked onto PY plates containing tetracycline, 0.0005% crystal violet and 0.02% 2,3,5-triphenyltetrazolium chloride. A single bioluminescent colony was then inoculated into PY medium containing 0.1% L-arabinose and cultured overnight at 28°C to induce transposition. The culture was streaked onto PY plates with crystal violet and 2,3,5-triphenyltetrazolium chloride and placed in an incubator at 30°C for 2 days until single colonies appeared. Bioluminescent colonies were patched onto PY plates with or without tetracycline to select tetracycline-sensitive colonies, confirming the absence of pBJ7. Chromosomal integration of *luxCDABE* was determined by sequencing.

### Deletion of *cmaA* from *Pto*-lux

Two DNA fragments containing the 1kb upstream or downstream region of *cmaA* were PCR-amplified from genomic DNAs of *Pto* using the primer sets cmaA_up1000_F plus cmaA_del_Rev or cmaA_del_Fwd plus cmaA_down_1000_Rev. These fragments were assembled into *Xba*I-digested pK18mobsacB. This construct was introduced into *Pto*-lux by electroporation. Bacteria were recovered in KB medium at 28°C for 2 hours and spotted onto a KB plate with rifampicin and kanamycin. The deletion of *cmaA* was induced by counter-selection on a KB plate containing rifampicin and 10% sucrose and confirmed by size reduction of a PCR fragment amplified by cmaA_conf_Fwd plus cmaA_conf_Rev. Primers used for generation of *Pto*-lux Δ*cmaA* are listed in Table S2.

### Inoculation of plants with *P. syringae*

Bacterial suspensions in sterile water were directly infiltrated using a needleless syringe into leaves of 4 to 5-week-old *A. thaliana*, 5 to 6-week-old *N. benthamiana*, or 3 to 4-week-old *S. lycopersicum*. In each independent experiment, one biological replicate consisted of 4 leaf discs (4 mm diameter), and 6-8 biological replicates were collected for each treatment. Alternatively, 4 to 5-week-old *A. thaliana* plants were spray-inoculated with bacterial suspensions in 10 mM MgCl_2_ with 0.04% Silwet-L77. After inoculation, plants were kept under covers for 1 hour and then the covers were removed. In each independent experiment, one biological replicate consisted of 6 leaf discs (4 mm diameter) excised from 3 different leaves of the same plants, and 8 biological replicates were collected for each treatment. *M. polymorpha* gemmae were placed on half-strength Gamborg’s B5 agar covered with a Whatman filter paper (Cat. No. 1001-085) and grown for 14 days at 22°C under continuous white LED light. The 14-day-old plants were transferred to fresh empty Petri dishes, submerged in the bacterial suspensions, and incubated with or without vacuum for 5 minutes. After the incubations, plants were placed onto a Whatman filter paper which had been soaked with Milli-Q water in fresh empty Petri dishes. Inoculated plants were kept in a climate chamber at 22°C under 16 h light/8 h dark cycle. Alternatively, *M. polymorpha* gemmae were grown on half-strength Gamborg’s B5 agar without the filter paper, and the bacterial suspension was poured into the cultured agar plate and vacuum infiltrated (Figure S7a). The bacterial suspension was decanted and plants were further cultured in a climate chamber at 22°C under 16 h light/8 h dark cycle. In each independent experiment, one biological replicate consisted of one thallus disc (5 mm diameter) excised from an individual plant, and 8 biological replicates were collected for each treatment.

### Quantification of *P. syringae* growth

Bioluminescence measurement was carried out with GloMax Navigator Microplate Luminometer (Promega) or Fluostar Omega (BMG Labtech) plate reader. For *A. thaliana, N. benthamiana*, and *S. lycopersicum*, lead discs (4 mm diameter) were punched out using a biopsy punch and placed in a white, light-reflecting 96 well plate (Corning, #3912). The plate containing leaf discs in each well was placed in the sample drawer of GloMax Navigator and kept in the dark by closing the lid for 10 min to reduce background signals. Then, luminescence was measured for 10 sec for each sample. After measurement, leaf discs were pulverized in 400 μl of 5 mM MgSO_4_ and serial dilutions were made. Of each dilution, 10 μl was streaked on KB plates containing rifampicin. Log_10_-transformed relative light units (RLU) and colony-forming units (CFU) per cm^2^ leaf surface area were calculated. For *M. polymorpha*, thallus disc (5 mm diameter) was punched from the basal region of thallus using a biopsy punch (Figure S7b). The thallus discs were transferred to wells of a white reflecting 96-well plate (VWR, #738-0016). The plate was placed in the Fluostar Omega (BMG Labtech) plate reader and kept in the dark for 10 min before measurement to reduce background signals. Then, luminescence was measured for 5 seconds for each sample. After measurement, thallus discs were ground in 400 μl of 10 mM MgSO_4_ using a mixing mill (MM400, Retsch) at 30 Hz for 5 min and serial dilutions were made. Of each dilution, 10 μl was streaked on KB plates containing rifampicin. The plates were then incubated at 22° C for 3 days until colonies became visible for counting. Log_10_-transformed relative luminescence units (RLU) and colony-forming units (CFU) per cm^2^ thallus surface area were calculated. Pairwise comparison was performed using the R function pairwise.t.test with pooled SD and the Benjamini-Hochberg method was used for correcting the multiple hypothesis testing. Linear regression and calculation of correlation coefficients were performed using the R function lm and cor, respectively.

### Bioluminescence imaging

Fusion SOLO S (VILBER) or ChemiDoc MP Imaging System (Bio-Rad) was used to visualize luminescence of plant tissues inoculated with *Pto*-lux. Detached leaves of *A. thalaiana* or *S. lycopersicum* were put on a wet paper and then placed in the sample drawer of the Fusion SOLO S. Before starting imaging, samples were kept inside the instrument with the lid closed to reduce background signals. *M. polymorpha* thalli on a Whatman filter paper were imaged using the ChemiDoc MP Imaging System. After 10 min of darkness adaptation, images were obtained using signal accumulation mode. The images were reduced to the signal showing pixels, hue shifted, and merged with microscopy images of the infected thalli using GNU image manipulation program.

## ACCESSION NUMBERS

The accession numbers for the genes discussed in this article are as follows: *RPM1* (AT3G07040), *RPS2* (AT4G26090), *DDE2* (AT5G42650), *EIN2* (AT5G03280), *PAD4* (AT3G52430), *SID2* (At1g74710), *AvrPto* (*AvrPto1*; PSPTO_4001), *AvrPtoB* (*hopAB2*; PSPTO_3087), *cmaA* (PSPTO_4709).

## ACKNOWLEDGEMENT

We thank Drs. Yuichi Takikawa (Shizuoka University) and Yasufumi Hikichi (Kochi University) for providing Japanese isolates of *P. syringae* and *R. solanacearum*, respectively, and Noriko Shimizu (Ritsumeikan University) for technical assistance. We also thank Drs. Kenichi Tsuda (Huazhong Agricultural University) and Tatsuya Nobori (Salk Institute) for critical reading of the manuscript. pBEN276, pBBR1MCS-2 and pFREE-RK2 were gifts from Pierre Germon (Addgene plasmid # 85168), Kenneth Peterson (Addgene plasmid # 85168) and Morten Norholm (Addgene plasmid # 92054), respectively. pK18mobsacB was obtained from National BioResource Project (NIG, Japan): E. coli. The tomato resource used in this study was provided by the National BioResource Project (NBRP), MEXT, JAPAN. Figures S2&S7 are created with BioRender.com. This work was supported by JST PRESTO (JPMJPR17Q6) and Grant-in-Aid for Scientific Research (B) (19H02960) to A.Mi, Ritsumeikan Global Innovation Research Organization to A.T., and supported by funds from the Max Planck Society and the ‘Priority Programme 2237 MAdLand’ funded by the Deutsche Forschungsgemeinschaft (NA 946/1-1) to H.N.

## AUTHOR CONTRIBUTIONS

A.Mi. designed the research; A.Mi. and H.N. conceived the experiments; A.Mi., A.Ma., T.S., K.M., and H.N. performed the experiments and analyzed the data; A.Mi., H.N., and A.T. wrote the paper.

---

**Figure S1.**
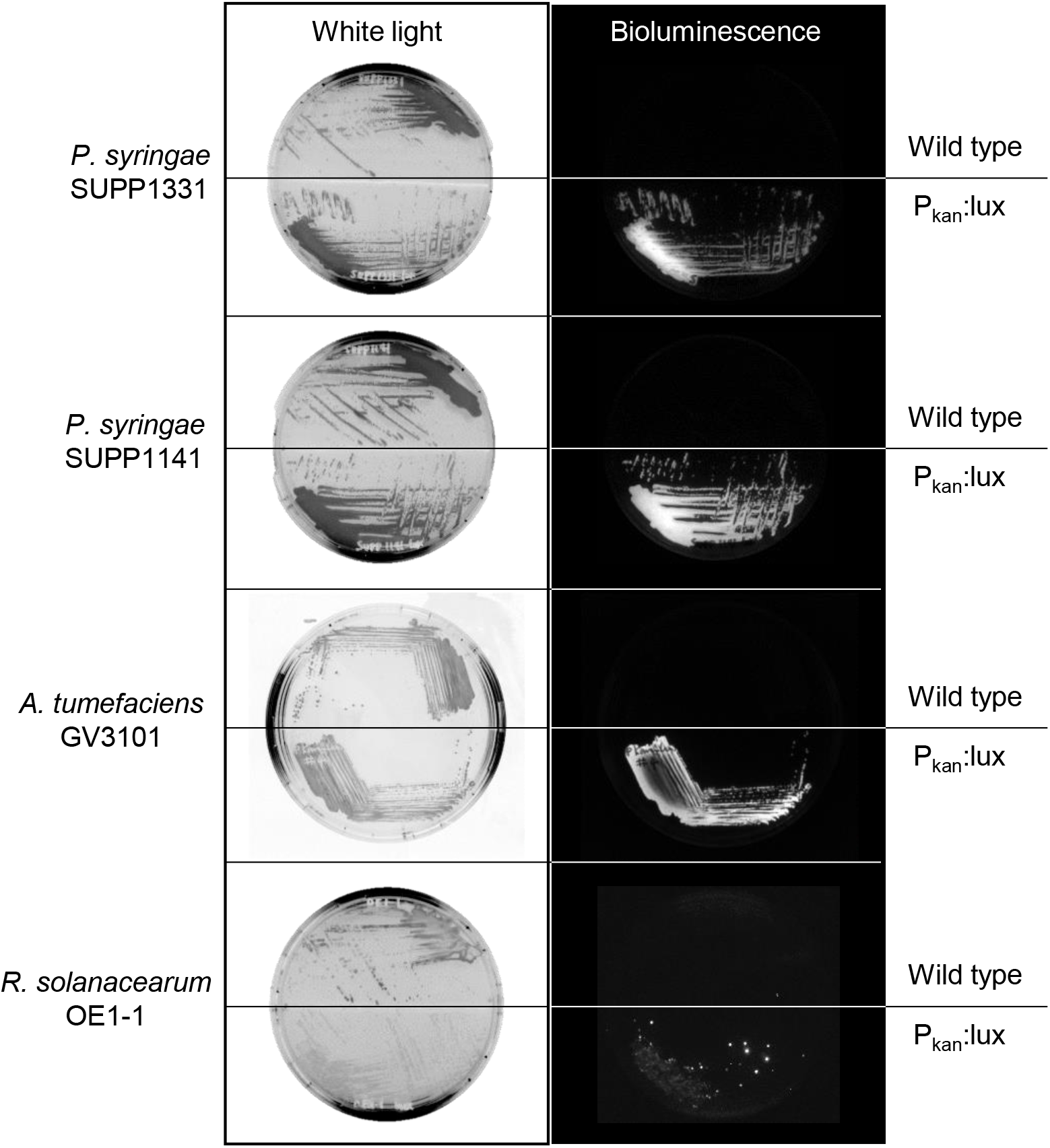
Generation of bioluminescent transformants of Japanese isolates of *P. syringae* SUPP1331 and SUPP1141, *Agrobacterium tumefaciens* GV3101 and *Ralstonia solanacearum* OE1-1. Wildtype and *luxCDABE*-tagged bacteria were observed under white light (left panels), and their bioluminescence was detected in the dark (right panels).

**Table S1.**
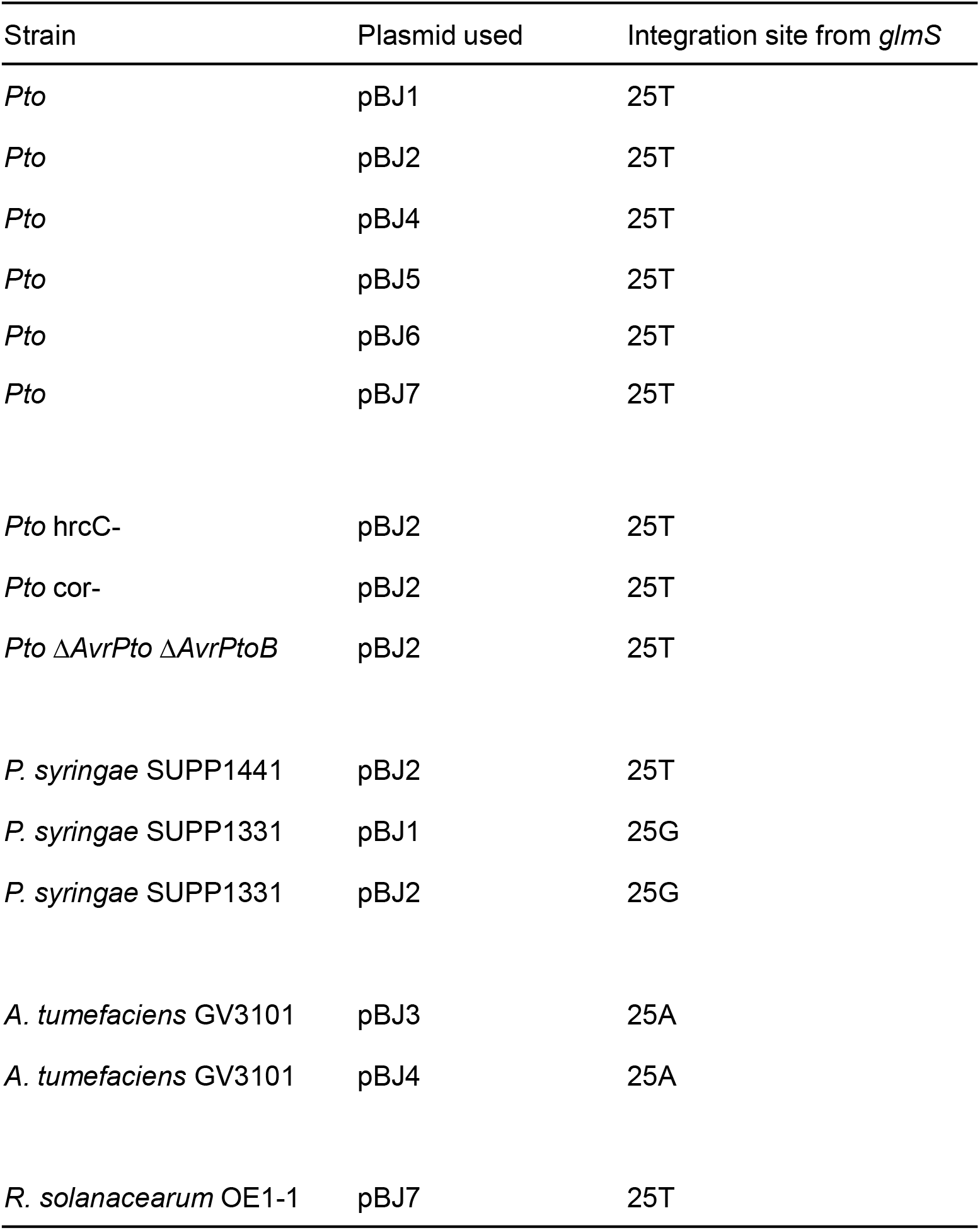
The positions of *luxCDABE* insertion in the genomes of various bacteria. Plasmids used to transpose *luxCDABE* are shown. Insertion of *luxCDABE* at attTn*7* downstream of *glmS* gene was determined by sequencing.

**Figure S2.**
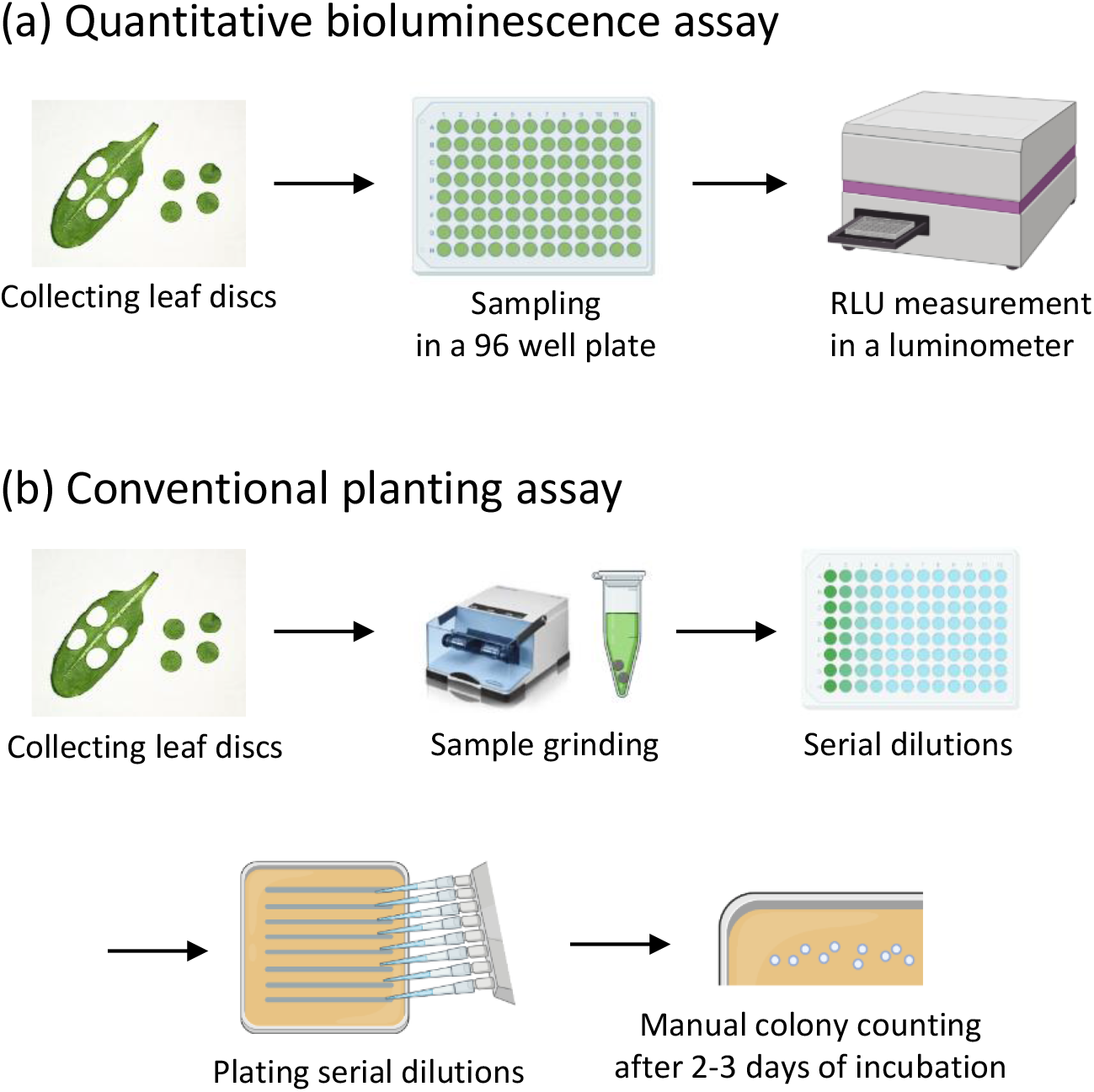
Comparison between quantitative bioluminescence assays and conventional colony counting assays. (a) In quantitative bioluminescence assays, a 96 well plate containing leaf discs in each well is placed in a luminometer and kept in the dark for 10 min. Then, bioluminescence intensities (RLU) are directly measured. (b) In conventional colony counting assays, samples are prepared by grinding leaf discs and are then used to make serial dilutions. These dilution series are streaked onto selective plates. After 2-3 days of incubation, bacterial colonies on the plates are manually counted.

**Figure S3.**
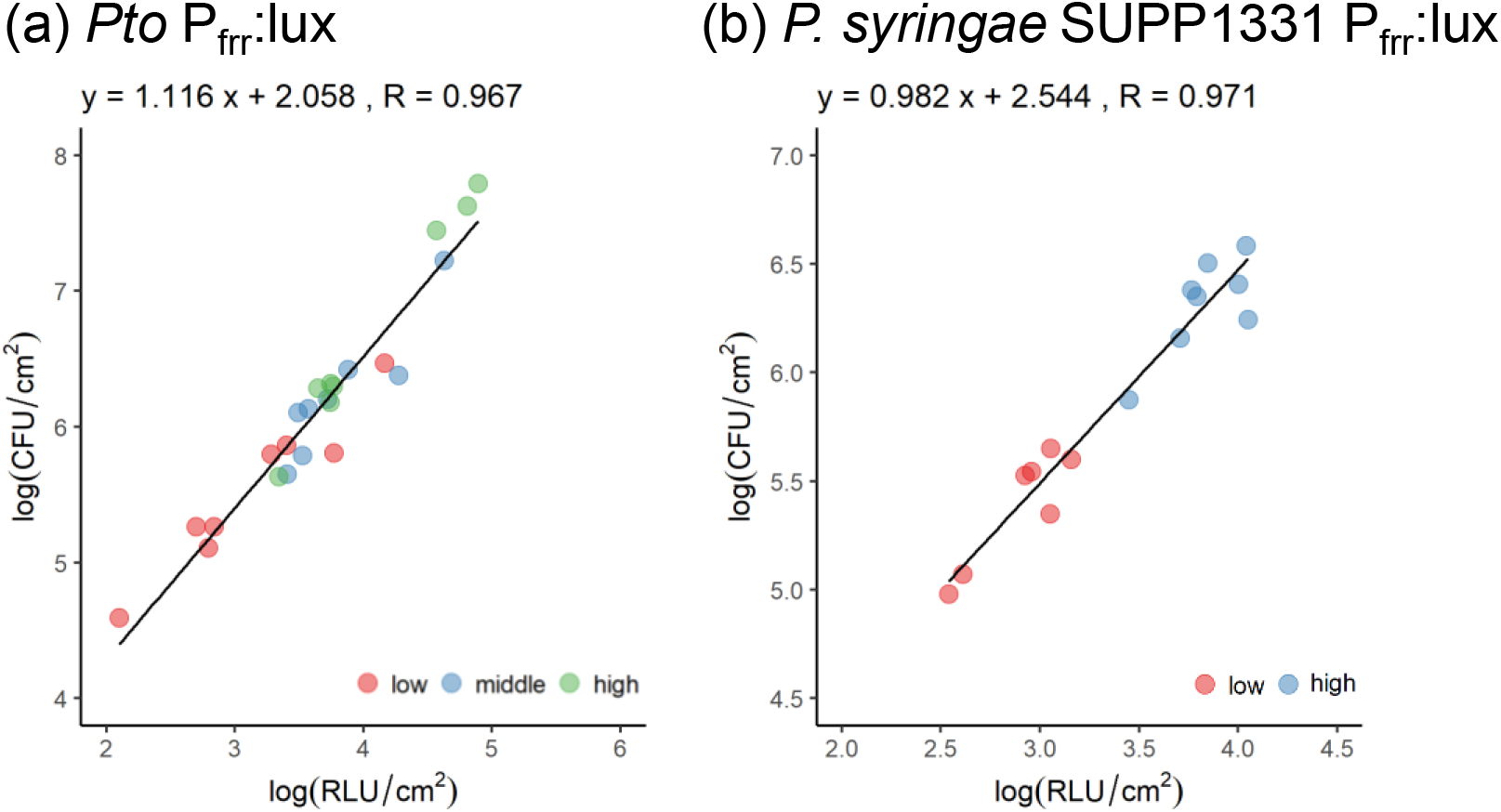
Correlations between CFU and RLU of *Pto* P_frr_:lux (a) and of *P. syringae SUPP1331* P_frr_:lux (b) in *A. thaliana*. (a and b) Leaves *of A. thaliana* were syringe-infiltrated with *Pto* P_frr_:lux (a) and *P. syringae* SUPP1331 P_frr_:lux (b) at varying doses (From low to high: OD_600_ = 0.0002, 0.001, 0.01 for *Pto* P_frr_:lux; 0.0002, 0.001 for and *P. syringae* SUPP1331 P_frr_:lux). Log_10_-transformed colony forming unit (CFU) and relative light unit (RLU) are plotted along with Y and X axes, respectively. Regression equations and Pearson’s correlation coefficients (R) are shown above the plots.

**Figure S4.**
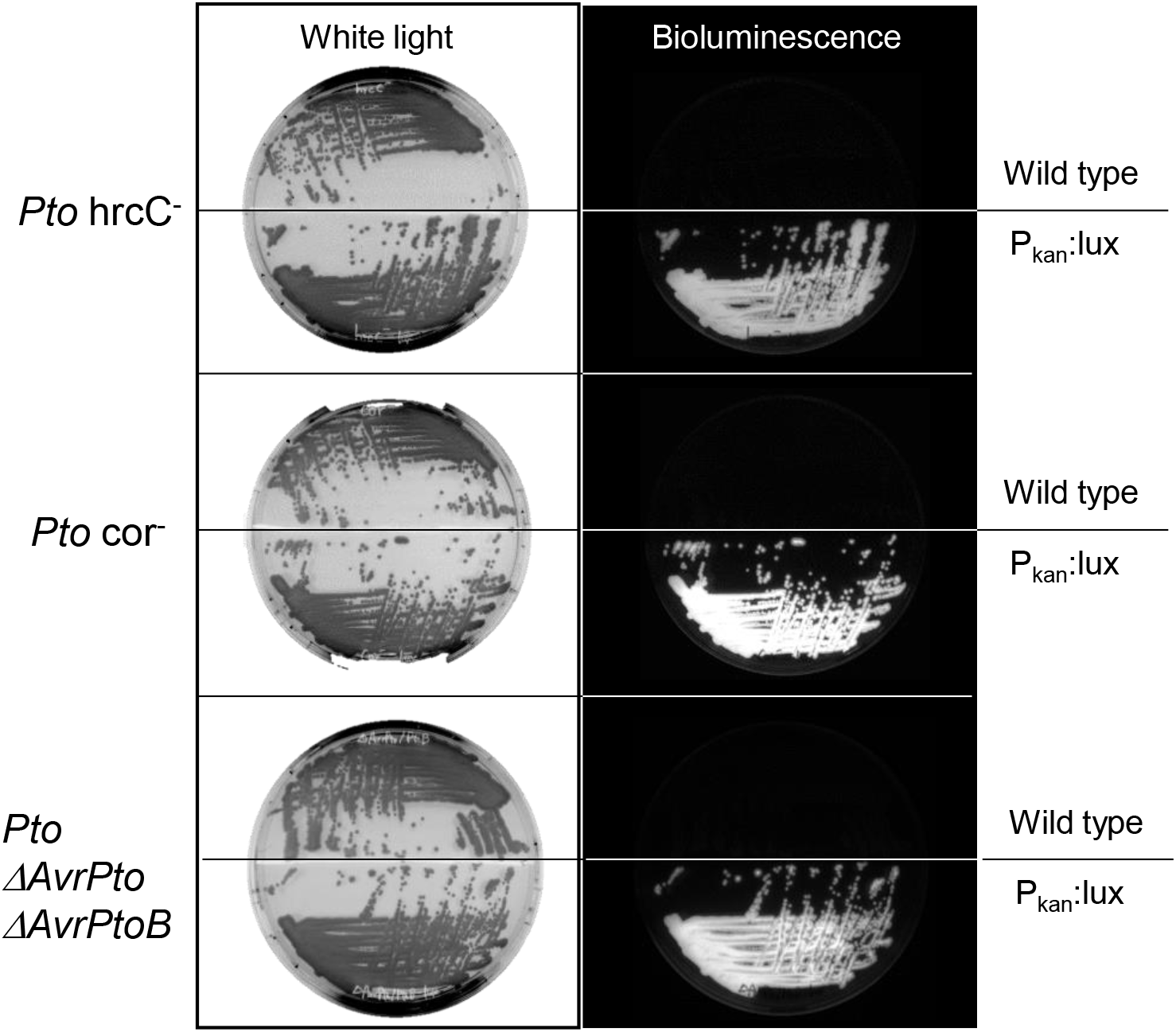
Generation of bioluminescent transformants of *Pto hrcC-, Pto cor- and Pto ΔAvrPtoΔ AvrPtoB*. Wildtype and *luxCDABE*-tagged bacteria were observed under white light (left panels), and their bioluminescence was detected in the dark (right panels).

**Figure S5.**
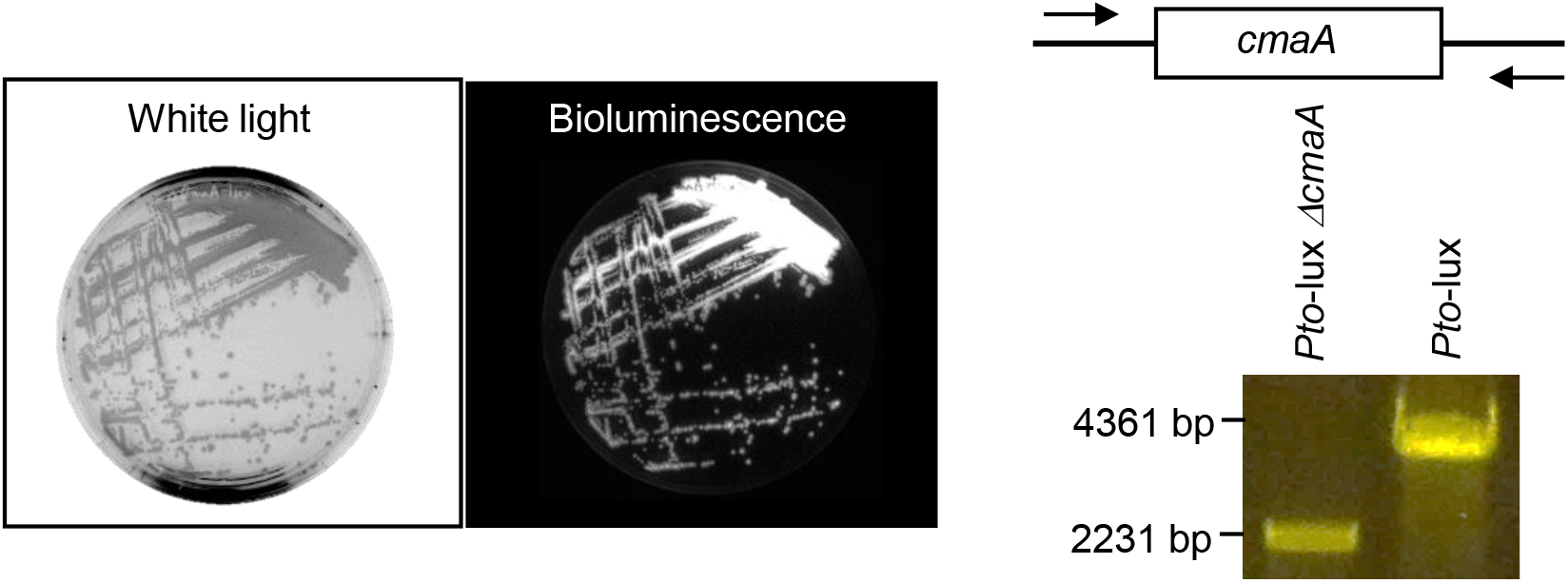
Deletion of *cmaA* from *Pto-*lux. (a and b) Unmarked deletion of *cmaA* was induced by homologous recombination using the suicide vector pK18mobsacB. (a) The resulting mutant, *Pto*-lux Δ*cmaA*, was observed under white light (left panel), and their bioluminescence was detected in the dark (right panel). (b) The deletion of *cmaA* was confirmed by size reduction of a PCR fragment spanning the gene.

**Figure S6.**
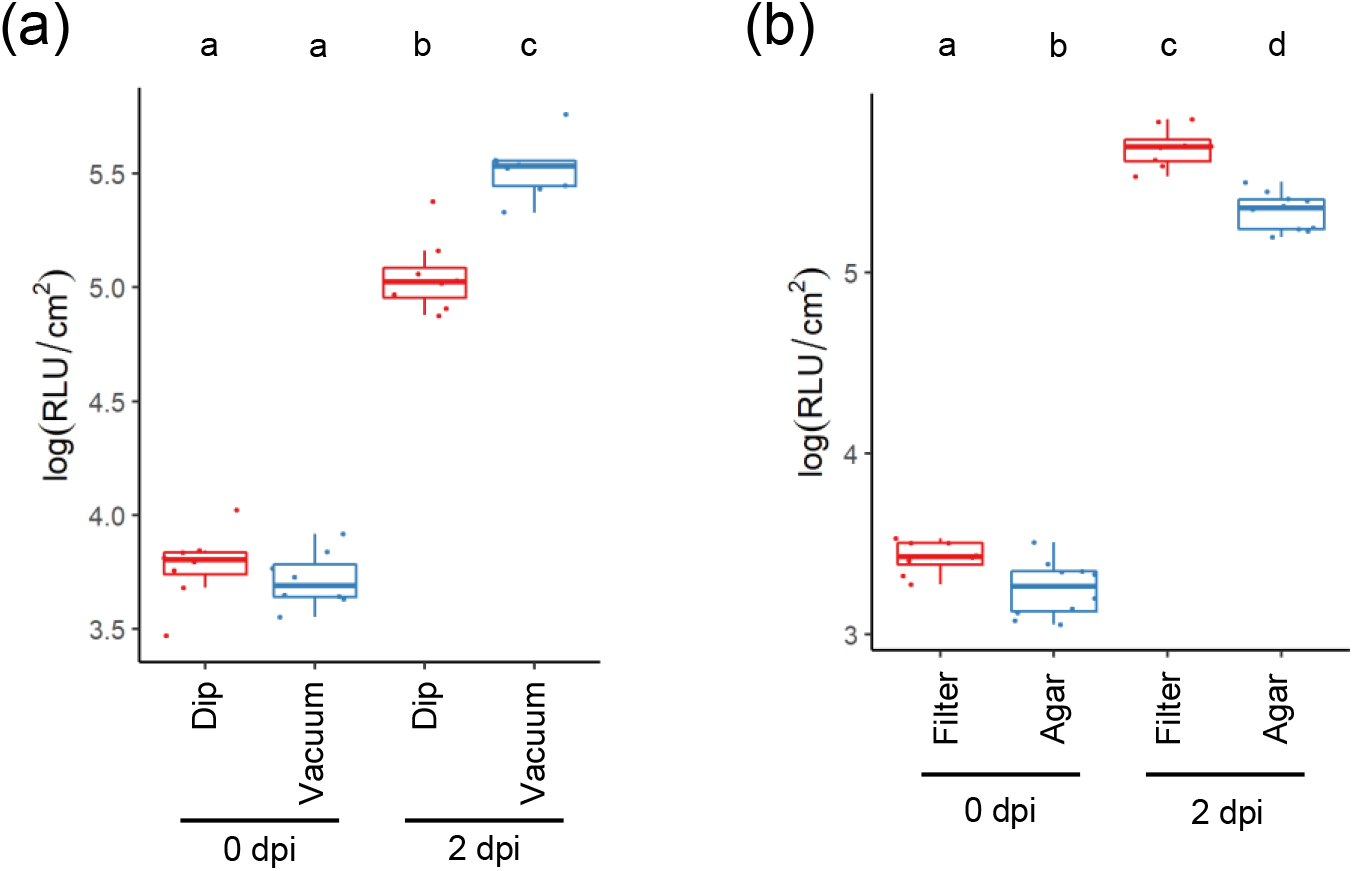
Inoculation methods affect growth of *Pto-*lux in Tak-1. (a) Fourteen-days-old Tak-1 thalli were vacuum infiltrated or dip inoculated with *Pto*-lux at OD_600_ = 0.01, and sampled 0 and 2 dpi. (b) Fourteen-days-old Tak-1 thalli grown on media with or without filter papers were vacuum infiltrated with *Pto*-lux at OD_600_ = 0.01, and sampled at 0 and 2 dpi. (a and b) Log_10_-transformed RLU were calculated from 8 biological replicates collected from thalli of different plants.

**Figure S7:**
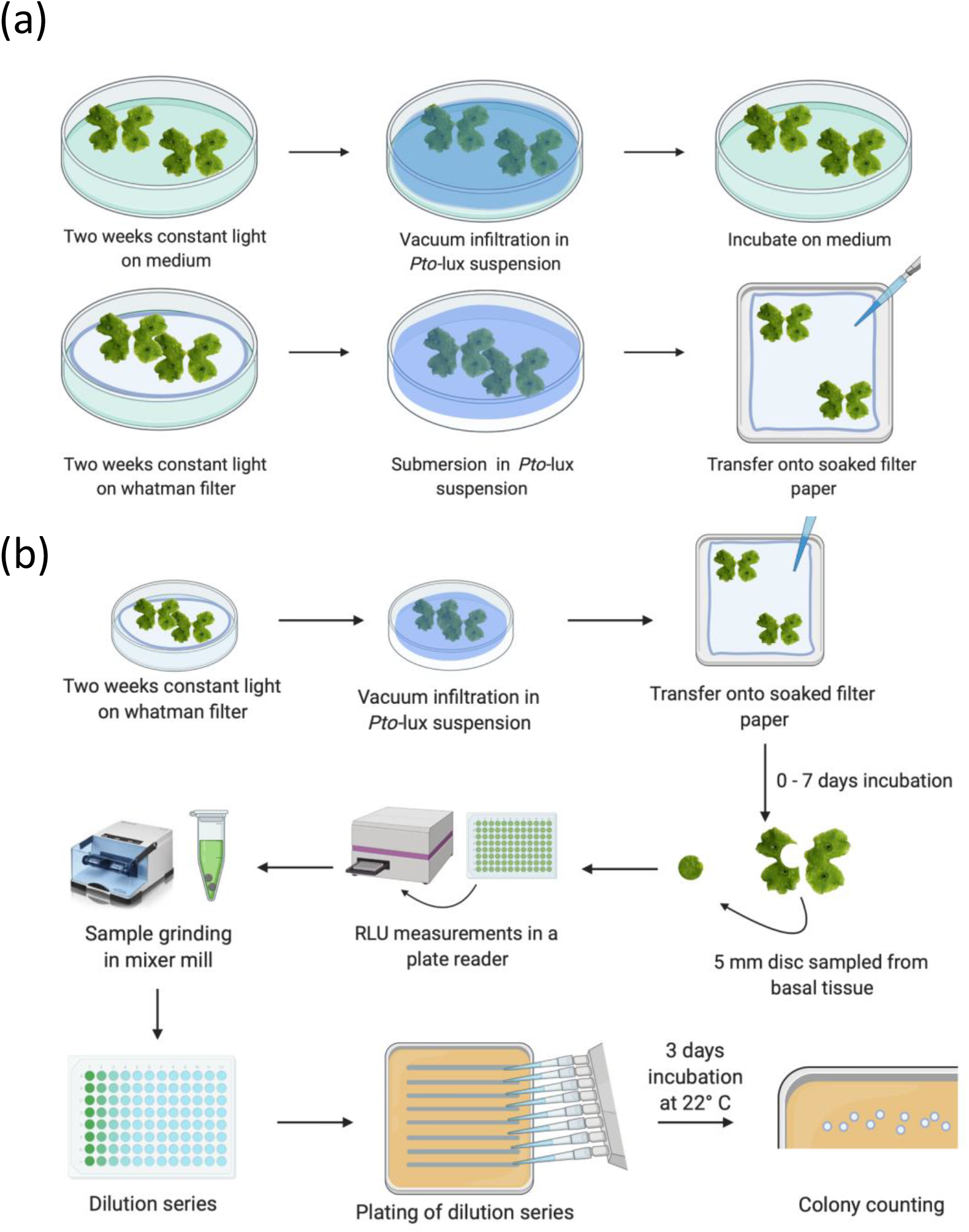
Graphical summary of the workflow of *Pto-*lux infection assays in *M. polymorpha*. (a) Plain submersion (dip) in the bacterial suspension compared to 5 min vacuum treatment while submerged (vac). (b) After vacuum infiltration with *Pto*-lux, thallus disc is excised and put into a well of a 96 well plate. The plate is placed in a luminometer and kept in the dark for 10 min. Then, bioluminescence intensities (RLU) are directly measured. After bioluminescence measurement, the same leaf discs were subjected to conventional colony counting assays to determine bacterial titers.

